# Genome-replicating HC-AdV – a novel high-capacity adenoviral vector class featuring enhanced *in situ* payload expression

**DOI:** 10.1101/2025.04.02.646678

**Authors:** Jonas Kolibius, Fabian Weiss, Patrick C. Freitag, Andreas Plückthun

## Abstract

High-capacity adenoviral (HC-AdV) vectors offer large transgene capacities and long-term expression of therapeutics, but require high doses due to limited transgene expression. In contrast, replication-competent AdV (RC-AdV) vectors enhance *in situ* transgene expression by genome replication and increased transcription from amplified genomes. Yet, RC-AdVs are constrained by minimal payload capacity, progeny formation, and toxic protein expression leading to rapid host cell death. To address these limitations, we developed a novel, genome-replicating HC-AdV vector. Therefore, we investigated AdV genome replication independently of progeny particle formation, and developed cell-based *trans*-replication assays, enabling us to probe the requirement for individual AdV proteins in AdV genome replication. We identified seven AdV proteins from the early transcriptional units which promote potent replication of HC-AdV genomes. We then created a genome-replicating HC-AdV vector by encoding an engineered minimal replication system that functionally reconstitutes AdV genome replication. Host cell transduction with our genome-replicating HC-AdV promoted *cis*-replication of the delivered HC-AdV genome and up to 20-fold increased reporter fluorescence. Our novel vector retained a large transgene capacity (22 kb) and, unlike RC-AdVs, did not induce a cytopathic effect nor host cell killing. Together, these data describe a novel delivery platform potentially allowing more efficacious vaccination and vector-mediated therapies.

## Introduction

Continuous advances in precision medicine have sparked a renaissance in the field of gene therapy, leading to the development of promising solutions for various indications such as cancer, monogenic, infectious, ophthalmological, neurological, and hematological diseases.^1–4^ *In vivo* gene therapeutics offer great potential, as they are directly administered, either intravenously (i.v.) or as targeted injection into the patient’s afflicted organ.^4,5^ Delivery of the therapeutic transgene to the target cell’s nucleus has been achieved using non-viral methods^6–8^ but for therapy predominantly by viral vectors,^2,3,9,10^ as reflected by their widespread use in 89% of gene therapies currently under development.^3^ The most commonly used viral vectors for *in vivo* gene delivery in clinical and preclinical applications are engineered and replication-deficient forms of the adeno-associated virus (AAV) and the adenovirus (AdV).^3,9^

AdVs feature a linear, double-stranded DNA genome, packed into a non-enveloped, icosahedral capsid of approximately 90-100 nm diameter. Over 200 non-human and >100 human AdV types (classified into species A to G) have been identified, with a considerable fraction thereof being vectorized.^11^ The human wild-type (WT) AdV-C5 genome has a length of ∼36 kb and encodes four early (E) (E1-4) and five late (L) (L1-5) transcriptional units (TUs). The early TUs are expressed prior to AdV genome replication, and the vast majority of the late TUs are expressed post-replication.^12^

AdV vectors were engineered in many aspects to exploit the AdV’s natural ability to transduce a diverse set of cell types and efficiently deliver the transgene to the nucleus.^13–15^ High-capacity AdV vectors (HC-AdVs) (also called “helper-dependent AdV vectors” (HD-AdVs), “gutless AdV vectors”, or “third-generation AdV vectors”) are particularly favorable, as they are fully devoid of any AdV protein-coding sequences. All genes are replaced with optimized stuffer DNA,^16^ and the HC-AdVs only contain two *cis*-acting elements, the AdV inverted terminal repeats (ITRs) and the packaging signal Ψ. Hence, HC-AdVs provide a large transgene capacity of up to 37 kb. During production, a co-transduced helper virus (HV) supplies all AdV proteins in *trans* which are required to express the capsid, *trans*-replicate the HC-AdV genome, and package it into viral particles. Their high transduction efficiency,^13,17,18^ modifiable tropism,^19–25^ stable gene expression,^11,14,26^ and ability to carry large transgenes of up to 37 kb^9,14,27^ make HC-AdVs particularly suitable for targeted DNA delivery of large and complex genetic circuits. Their vector genomes remain and persist in the nucleus as episomes,^28^ posing only a minor risk of insertional oncogenesis. HC-AdV genome association to cellular histones can promote stable and long-term expression of the encoded transgenes for up to seven years,^29–31^ although a certain reduction in transgene expression has been observed over time, most likely due to physiological cell turnover and the accompanying loss of the extrachromosomal vector genome.^29^ These attributes are especially advantageous when addressing complex and multifaceted diseases, such as cancer. As previously shown, AdV vector-mediated immunotherapy allows sustained combinatorial *in situ* expression^32,33^ of synergistically acting therapeutic proteins, enhancing their efficacy in such challenging therapeutic contexts.^32^

Despite all favorable features, *in vivo* administration of HC-AdV vectors is still limited by two major aspects, which currently compromise broader application of HC-AdV-based gene therapies. First, limited payload expression levels require high vector doses, and second, high vector doses elicit potent innate immune responses.^3,4^ This highlights a central problem of HC-AdV vectors: since the acute toxicity is a direct consequence of the dose-dependent activation of the innate immune system by the vector, reduction of the administered vector dose is inevitable for safe therapies.^34^ Conversely, administration of decreased vector titers results in diminished tissue transduction, low expression of the therapeutic payload and therefore reduced efficacy.

A promising strategy to achieve clinical efficacy with a reduced HC-AdV vector dose is to enhance *in situ* expression of the AdV-encoded payload in the transduced host cell via AdV genome replication. During the WT AdV life cycle, genome amplification causes a drastic increase in late gene expression, promoting efficient progeny particle formation. AdV genome replication results in >100,000-fold amplification of genome copies.^12^ This can lead to more than 1000-fold increase in late gene transcription and substantially enhanced *in situ* expression of capsid proteins.^35,36^ With the development of replication-competent AdV (RC-AdV)^37,38^ and single-cycle replicating AdV (SC-AdV)^39,40^ vectors it was attempted to take advantage of this replication-dependent boost of transgene expression. RC-AdVs (e.g., conditionally-replicating and oncolytic AdV (OAdV) vectors) are characterized by completion of the entire AdV life cycle, including AdV genome replication, formation of progeny particles and cell lysis of the host cell. In contrast, SC-AdVs feature a deletion of an AdV gene, critical for particle formation or viral processing (e.g., protein IIIa,^41^ fiber,^42^ or protease^43^), and thus the viral life cycle is aborted after AdV genome replication, preventing the final step of particle assembly and progeny release.^40,41,44^ Neither RC-AdV nor SC-AdV are suitable for co-expression of multiple payloads due to their limited transgene capacities (<3 kb and <4.5 kb, respectively), and additionally, they exhibit a highly cytotoxic profile due to the expression of toxic AdV proteins, such as adenovirus death protein (ADP), protease, and E4 ORF1 and 4.^45–51^ Induction of this cytopathic effect causes rapid apoptotic clearance of host cells,^40^ rendering RC-AdVs and SC-AdVs unsuitable for sustained and long-term gene therapy applications, despite their high payload expression levels.

We hypothesized that it might be possible to fully uncouple AdV genome replication from late gene expression and therefore generate a new AdV vector class that features enhanced transgene expression as a consequence of genome amplification, yet without the cytotoxic effects observed for RC-AdVs and SC-AdVs. Here, we present the development of such a new HC-AdV vector class, which features self-induced and self-sustained genome replication within the transduced cell, indeed leading to increased payload expression. To develop such a system, we first established cellular *trans*-replication assays that allow screening of any AdV gene product for its contribution towards AdV genome replication. Using these assays, we identified in total seven AdV genes, which are required for AdV genome amplification.

Previous work has already established that all three E2 proteins, DNA binding protein (DBP), precursor terminal protein (pTP), and AdV DNA Polymerase (Ad Pol), are essential for *in vitro* AdV genome replication, as they reconstitute the actual viral DNA replication machinery.^12,52,53^ Briefly, AdV DNA replication is initiated by association of pTP with Ad Pol, and covalent addition of a deoxycytidine monophosphate (dCMP) to pTP. This complex recognizes the AdV core origin of replication, encoded by the AdV ITRs, and starts protein-primed DNA synthesis by annealing to nucleotide 4 (3’-GTA<u>G</u>TAGTTA) at the 3’-end of the ITR, while DBP concurrently binds to the displaced 5’-end.^54,55^ Upon synthesis of the first trinucleotide, CAT, the Ad Pol-pTP-CAT intermediate jumps back three nucleotides and hybridizes to bases 1-3 (3’-<u>GTA</u>GTAGTTA). This induces dissociation of Ad Pol from the pTP protein primer and thus increases its processivity. DBP binds cooperatively to the non-template single strand and provides the driving force for further ATP-independent unwinding of the double-stranded viral DNA through polymerization, enabling template-strand elongation and formation of a new duplex genome. The ITRs of the displaced single strand then hybridize either intra- or intermolecularly to restore functional templates for further genome amplification.^12,56^

Using cellular assays, we confirm here that all E2 proteins are essential for AdV replication, as expected, and we describe the optimal combination of further AdV proteins required for most efficient AdV replication in cells. Our developed *trans*-replication assays indicate that the combination of the three AdV E2 proteins, the three E1 proteins (E1A, E1B-55k, and E1B-19k) and the gene product of E4 ORF6 is required to promote potent AdV genome replication in cells. We present the design of an engineered, compact minimal AdV replication system that reconstitutes the natural AdV DNA replication machinery to rapidly amplify HC-AdV genomes. Most importantly, we demonstrate that transduction of E1-complementing and non-complementing cells with our genome-replicating HC-AdV results in robust *cis*-acting HC-AdV genome amplification and enhanced payload expression of the encoded reporter. Furthermore, we show that all AdV genes unrelated to replication can be excluded from the genome of our new HC-AdV vector, which thereby retains a large transgene capacity of >22 kb, potentially allowing potent *in situ* expression of multiple therapeutics.

## Results

### Development and validation of *trans*-replication assay to quantify AdV genome replication

The development of a genome-replicating HC-AdV first requires identifying the essential AdV gene products involved in DNA replication and those modulating the host cell environment to support this resource-consuming process. Afterwards, those essential AdV genes will be encoded on a HC-AdV vector to reconstitute a functional replication system in the transduced target cell, which should result in amplification of the HC-AdV genome in *cis* and increased transcription of vector-encoded genes, promoting enhanced payload expression (Figure 1A). Given the complexity of the AdV-C5 genome, which encodes more than 40 genes (Figure S1), we employed two complementary strategies: a top-down approach involving sequential deletion of AdV genes or TUs from a functional replication system, and a bottom-up approach to reconstruct a rationally designed, minimal replication system.

**Figure 1.**
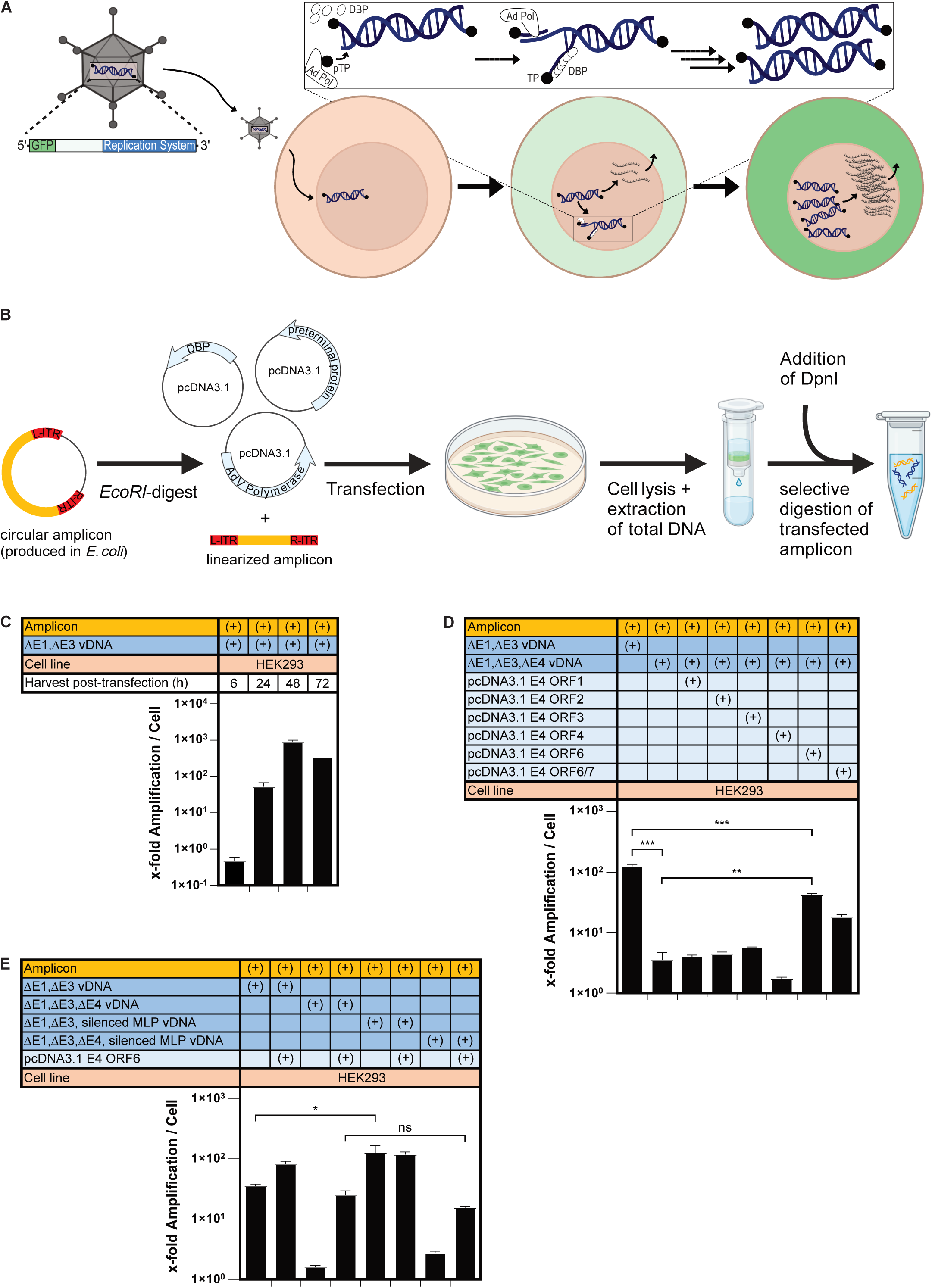
Identification of AdV protein components required for genome-replicating HC-AdV by *trans*-replication of AdV mini-chromosomes. (A) Conceptual representation of genome-replicating HC-AdV. Transduction of the host cell with genome-replicating HC-AdV initiates nuclear translocation of vector genome, followed by transcription of encoded replication machinery (including DBP, pTP, and Ad Pol) and payload (GFP). Prior to onset of replication, only low amounts of GFP are expressed due to a limited number of vector genomes that can serve as templates for transcription. Expressed components of AdV DNA replication machinery drive potent, protein-primed *in situ* replication of vector genomes (see magnified illustration), which promotes increased GFP transcription and expression without the formation of viral progeny particles. (B) Graphical depiction of *trans*-replication of an AdV mini-chromosome in HEK293 cells. When transfected plasmid(s) (light blue) express a functional AdV replication machinery, the *EcoRI*-linearized amplicon (orange) is *trans*-amplified upon recognition of accessible AdV ITRs (red). HEK293 cells are harvested and total DNA is extracted, *DpnI*-digested to remove methylated templates and subjected to qPCR measurement. (C) Quantification of x-fold replication per cell of the amplicon by qPCR, determined 6, 24, 48, and 72 h after co-transfection of amplicon with plasmids encoding either a functional (here ΔE1,ΔE3 vDNA) AdV replication machinery or non-functional (pC4HSU vDNA) machinery, with the latter used for normalization. X-fold amplification was analyzed by determining *DpnI*-digested amplicon copy numbers resulting from co-transfection with DNA to be probed for functional replication, relative to co-transfection with non-coding pC4HSU vDNA. (D) Determination of x-fold amplification of AdV mini-chromosome by qPCR, as analyzed 48 h post co-transfection of HEK293 cells with amplicon, ΔE1,ΔE3,ΔE4 vDNA, and individual E4 genes. (E) Quantification of x-fold *trans*-amplification via qPCR, 48 h after co-transfection of HEK293 cells with amplicon, ΔE1,ΔE3 vDNA, or ΔE1,ΔE3,ΔE4 vDNA (with each of the vDNAs additionally encoding a silenced major late promoter (MLP)), and E4 ORF6. Statistics: Representative data of two or three independent experiments are shown. Bar graphs represent mean x-fold amplification ± SD, n = 3. Statistical significance was determined by one-way ANOVA with Dunnett’s test for multiple comparisons. Not significant (ns) P > 0.05; *P ≤ 0.05; **P ≤ 0.01; ***P ≤ 0.001; ****P ≤ 0.0001.

Construction of our new vector class first required the dissection of AdV genome replication and progeny formation. AdV protein functions were traditionally studied by generating AdV virions containing the corresponding gene knockouts. Hence, previous research has largely examined the functions of certain AdV proteins across the full AdV life cycle rather than scrutinizing their specific roles in discrete processes, such as genome replication or transcription from *de* novo synthesized genomes. However, it is technically challenging and often not feasible to generate AdV vectors with deletions of entire TUs (particularly the late gene TU with its introns) or, alternatively, to create HC-AdV vectors encoding an individual set of viral genes. To circumvent these limitations we developed a transfection-based assay to screen for AdV gene products required for genome replication in cells. Since quantifying *cis*-replication is limited to testing ITR-containing AdV genomes, we instead determined *trans*-replication of an AdV mini-chromosome (amplicon), as described by Hay *et al*.^57^ This approach enabled us to precisely determine the contribution of individual AdV proteins to genome replication, while effectively eliminating crosstalk and interactions with other AdV proteins. Quantification of resulting changes in amplicon copy numbers via qPCR (Figure 1B) allowed systematic investigation of specific AdV gene combinations and their capability to reconstitute functional AdV genome replication.

The AdV “mini-chromosome” containing the left and right AdV ITRs was cloned and propagated as a plasmid in *E. coli*. The plasmid was linearized to release the ITRs, which serve as origin of AdV replication, and co-transfected into AdV E1-complementing HEK293 cells along with DNA encoding a functional AdV replication machinery (e.g., ΔE1,ΔE3 viral DNA (vDNA)). To clearly distinguish between transfected and *de novo* synthesized amplicons, the extracted total DNA was digested with *DpnI*, which selectively cleaves and depletes bacterially methylated template DNA^58^ and thus prevents qPCR detection of transfected amplicon (Figure S2A). This approach allowed precise quantification of replication efficiency, as only newly synthesized (non-methylated) amplicon DNA served as template during the qPCR reaction.

To validate our cellular *trans*-complementation assay, we first co-transfected HEK293 cells with linearized amplicon and either non-coding pC4HSU^16^ vDNA or ΔE1,ΔE3 vDNA encoding a functional replication machinery. At 6 h post-transfection (p.-t.), no difference was detectable in sensitivity of the extracted amplicon towards *DpnI*-digestion when co-transfected with either vDNA (Figure S2B). However, 24 h p.-t., qPCR quantification of the amplicon suggested a partial resistance of amplicon DNA towards *DpnI*-digestion when co-transfected with a functional replication machinery, such as encoded by ΔE1,ΔE3 vDNA. Forty-eight and 72 hours p.-t., amplicons co-transfected with ΔE1,ΔE3 vDNA showed full resistance to *DpnI*-digestion, while those with non-coding vDNA remained sensitive. This indicates that *DpnI*-resistance is indeed a direct consequence of a functional AdV replication machinery being present, promoting amplicon *trans*-replication in a mammalian cell. When *DpnI*-digested amplicon copies were quantified per cell (using GAPDH as qPCR reference gene), *trans*-replication was indicated as x-fold amplification upon normalization to amplicon copy numbers after co-transfection with non-coding pC4HSU vDNA (Figure 1C). Co-transfection with ΔE1,ΔE3 vDNA resulted in 100 to 1,000-fold replication of the amplicon.

We then were curious about the role of the individual AdV gene products from the E4 TU in supporting AdV genome replication in cells. The E4 TU encodes six individual proteins (E4 ORF1, ORF2, ORF3, ORF4, ORF6, and ORF6/7), and deletion of the whole E4 TU from a RC-AdV results in loss of progeny formation and replication-deficiency,^59^ which can be individually rescued by E4 ORF3 or ORF6.^60,61^ However, the multifunctional roles of E4 ORF3 and ORF6 had previously only been examined in the context of an otherwise WT infection, with a focus on progeny formation, rather than genome replication.

E4 ORF3 primarily disrupts host defense mechanisms like DNA damage repair (DDR) and p53 transcriptional activities through reorganization of promyelocytic leukemia nuclear bodies (PML-NB).^62–65^ In contrast, E4 ORF6 plays a broader role in replication, mRNA processing and export, and host protein degradation, by co-opting a cellular ubiquitin ligase complex (consisting of cullin 5 (Cul5), Rbx1, and elongins B and C) to modify the cellular environment for efficient viral replication.^66–70^ Both E4 ORF3 and ORF6 require interaction with the E1B-55k gene product to exhibit their full functional capacities, and they are functionally redundant in restoring the AdV life cycle of ΔE4 AdV-C5, with E4 ORF6 being more efficient in promoting replication and progeny formation.^61^ To determine the role of the different E4 gene products specifically in the context of AdV genome replication (and not the whole life cycle), HEK293 cells were co-transfected with linearized amplicons and either ΔE1,ΔE3 vDNA or ΔE1,ΔE3,ΔE4 vDNA, along with plasmids encoding the individual E4 open reading frames (ORFs). Deletion of the E4 TU caused a near-complete loss of AdV replication, evident from the substantial reduction in amplicon replication per cell (Figure 1D). In line with previous reports, this defect was fully rescued by *trans*-complementation with E4 ORF6, highlighting the critical role of the E4 ORF6 gene product in AdV genome replication. Partial rescue was observed upon co-transfection with an expression plasmid encoding the fusion protein E4 ORF6/7, likely due to increased E2 gene expression and altered E2F-4 nuclear localization.^71,72^ In contrast, plasmids expressing E4 ORF1, 2, 3, or 4 failed to restore replication, indicating their minimal contribution towards reconstituting the AdV replication machinery, which is particularly interesting in regard to E4 ORF3.

Expression of most AdV late genes is regulated by the major late promoter (MLP), which remains at basal activity during early stages of infection and is fully activated only after genome replication, resulting in expression of most AdV late genes after genome amplification. To further scrutinize the role of late genes in replication, we generated ΔE1,ΔE3 and ΔE1,ΔE3,ΔE4 vDNAs with multiple mutations in the MLP transcription factor binding sites (TATA box, upstream element (UPE), and inverted CAAT box) (Figure S3A),^35,73–76^ thereby silencing the promoter while preserving the coding sequence for AdV Polymerase on the antisense strand (Figure S1A and S3A). Western blotting and immunostaining of three representative late genes from the L1, L4 and L5 TUs confirmed effective MLP silencing (Figure S3B). We then assessed *trans*-replication of our amplicon by the four different vDNAs. ΔE1,ΔE3 vDNA promoted similar levels of replication regardless of the activity of the MLP (active or silenced), suggesting that AdV genome replication does not depend on the presence of AdV late gene products (Figure 1E). This finding was confirmed when we monitored *trans*-replication of the amplicon by ΔE1,ΔE3,ΔE4 vDNA and ΔE1,ΔE3,ΔE4, silenced MLP vDNA. The amplicon was only replicated by the transfected machinery when E4 ORF6 was *trans*-complemented, however replication occurred independently of the functionality of the MLP. Similar observations were made in the alternative E1-complementing cell line 911,^77^ further corroborating our findings (Figure S4).

In summary, we developed a *trans*-replication assay to quantify AdV genome replication in cells and identified viral gene products involved in DNA replication in the cellular environment. Using this assay, we showed that deleting the E4 transcription unit abolishes replication, which can be fully rescued only by *trans*-complementation with E4 ORF6. We conclusively demonstrated that late genes are indeed not required for AdV DNA replication to occur.

### Seven AdV proteins encoded by E1, E2 and E4 ORF6 are sufficient for replication of an AdV mini-chromosome in an E1-non-complementing cell line

Given the well-established critical role of DBP, pTP, and Ad Pol in *in vitro* replication assays,^53^ we next pursued our bottom-up approach and co-transfected HEK293 cells with the amplicon and expression plasmids encoding DBP, pTP, Ad Pol, and E4 ORF6 (Figure 2A). These four proteins proved to be sufficient to drive over 200-fold replication of the amplicon, exhibiting a similar degree of replication efficiency as observed for the positive controls ΔE1,ΔE3 vDNA and ΔE1,ΔE3,ΔE4 vDNA + E4 ORF6 in E1-complementing HEK293 cells. We hence concluded that the remaining AdV genes encoded on ΔE1,ΔE3 vDNA are unlikely to be mechanistically involved in AdV DNA replication, nor do they support replication by altering the host cell environment.

**Figure 2.**
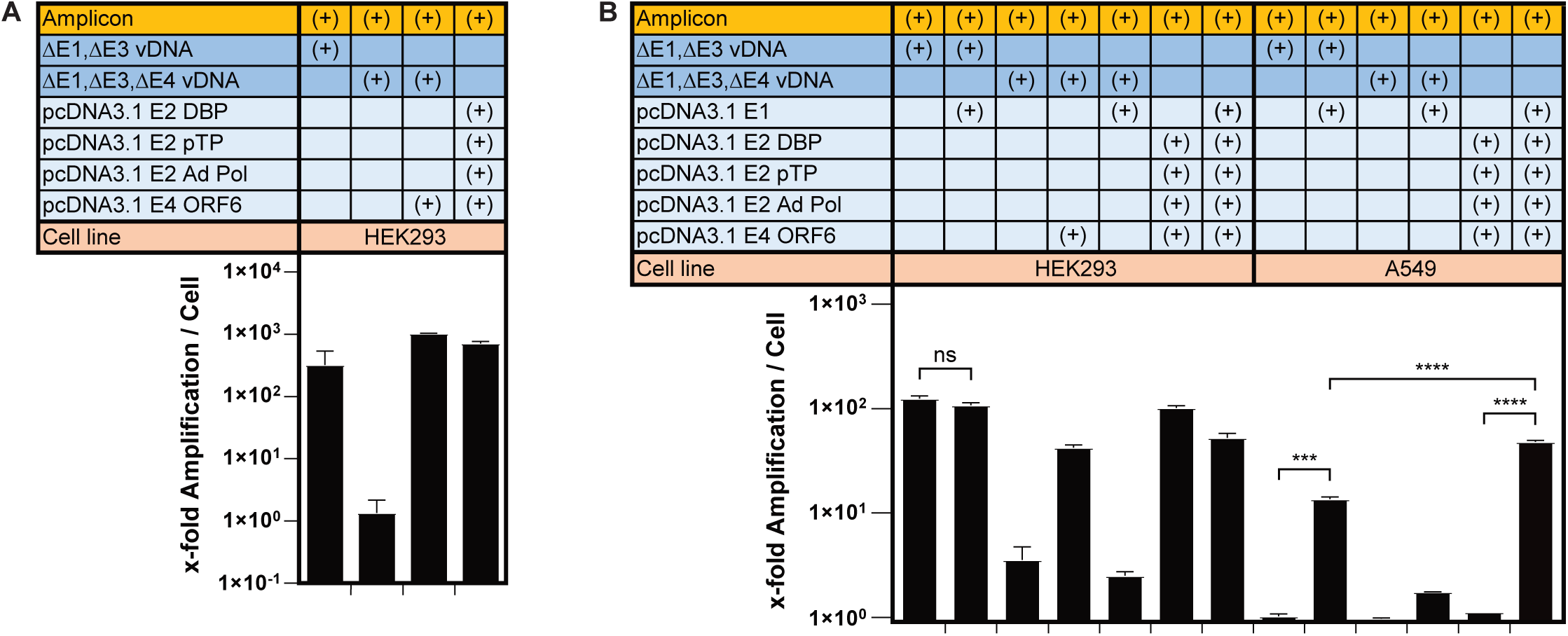
E1, E2 and E4 ORF6 gene products promote *trans*-replication of AdV mini-chromosome in E1-non-complementing A549 cells. (A) qPCR analysis of x-fold amplification of AdV mini-chromosome (yellow) 48 h after co-transfection of HEK293 cells with different vDNAs (dark blue) and individual expression plasmids (light blue). Cells were transfected with control viral DNA (ΔE1,ΔE3 vDNA or ΔE1,ΔE3,ΔE4 vDNA) and expression plasmids encoding AdV DBP, pTP, Ad Pol, or E4 ORF6. (B) qPCR quantification of x-fold amplification of mini-chromosome (yellow) upon co-transfection of E1-complementing HEK293, and E1-non-complementing A549 cells. Cells were co-transfected with control replication systems (ΔE1,ΔE3 vDNA, and ΔE1,ΔE3,ΔE4 vDNA; dark blue) and combinations of expression plasmids (light blue) encoding for the entire E1 TU (including E1A, E1B-19k, and E1B-55k), DBP, pTP, Ad Pol, or E4 ORF6 and harvested 48 h post-transfection for qPCR analysis. Statistics: Representative data of two or three independent are experiments shown. Bar graphs represent mean x-fold amplification ± SD, n = 3. Statistical significance was determined by one-way ANOVA with Dunnett’s test for multiple comparisons. Not significant (ns) P > 0.05; *P ≤ 0.05; **P ≤ 0.01; ***P ≤ 0.001; ****P ≤ 0.0001.

While E1 gene products (E1A, E1B-19k, and E1B-55k) are known for their cell-transforming properties,^78^ it was critical to further elucidate their specific role in AdV genome replication. AdV-C5 expresses five E1A isoforms that carry out multiple essential functions, including activation of viral early gene and host gene transcription, host chromatin reorganization and cell cycle progression.^35,78^ Importantly, E1A sequesters and inactivates tumor suppressors of the Retinoblastoma (Rb) gene family and thereby releases and activates transcription factors of the E2F family, promoting cell cycle progression from G0/G1 to S phase. This results in host cell transformation and transactivation of p53-responsive genes.^79^ The E1B TU encodes two proteins, E1B-19k and −55k, both mainly antagonize apoptotic signals induced by E1A. 19k blocks p53-independent apoptotic signals such as TNFα-mediated and Fas ligand-mediated death. E1B-55k is a multifunctional protein pivotal for inhibiting p53- and Daxx-mediated cell death, while also actively modifying the cellular environment to support efficient AdV genome replication and AdV gene expression via binding to other AdV proteins, such as E4 ORF6.^67,80^

Replication efficiency following co-transfection of HEK293 cells with the amplicon and ΔE1,ΔE3 vDNA was not enhanced by additional E1 expression (Figure 2B), indicating that sufficient levels of E1 protein are generally present in HEK293 cells for maximal AdV genome replication efficiency. The absence of an effect from further E1 overexpression in HEK293 cells was also confirmed by co-transfection of the amplicon in combination with ΔE1,ΔE3,ΔE4 vDNA and of the amplicon with E2 and E4 ORF6. As all AdV E1 genes are constitutively expressed in HEK293 cells, we next performed *trans*-complementation assays in the E1-non-complementing cell line A549 to examine the role of the AdV E1 proteins in the AdV genome replication machinery. Co-transfection of A549 cells with the amplicon and ΔE1,ΔE3 vDNA demonstrated that amplicon replication only occurs in the presence of a plasmid suppling all three E1 proteins. However, it remained unclear whether the E1 proteins are truly crucial for replication, or rather if E1A is primarily needed for the *trans*-activation of the E2 and E4 promoters encoded on ΔE1, ΔE3 vDNA, which is necessary for producing the components required for the replication machinery. Moreover, *trans*-replication with ΔE1,ΔE3,ΔE4 vDNA still required E4 ORF6, and E1 did not compensate for the lack of E4 ORF6. To clearly determine the role of E1 in AdV replication, we thus co-transfected A549 cells with a combination of different expression plasmids encoding DBP, pTP, Ad Pol and E4 ORF6 (all driving gene expression under the control of a constitutively active CMV promoter) with or without an additional expression plasmid encoding the E1 genes. While we could not detect amplicon replication in the absence of E1, we could measure a rescue of amplicon replication only when E1 was present.

In summary, we identified seven proteins (E1A, E1B-19k, E1B-55k, DBP, pTP, Ad Pol, E4 ORF6) as essential for successful replication of an AdV mini-chromosome in E1-non-complementing cells. E1 is therefore indispensable not only for *trans*-activating E2 and E4 promoters for expression of the replication machinery but also for modifying the cellular environment to support the demanding process of AdV DNA replication. Other AdV genes do not play a significant role in genome replication.

### E1, E2 and E4 ORF6 are sufficient to replicate native, incoming HC-AdV genomes in the E1-non-complementing cell line A549

Despite having gained valuable insights from *trans*-amplification, quantifying AdV genome replication using an AdV mini-chromosome has two major caveats. First, unlike natural AdV genomes, the transfected amplicon is not delivered to the nucleus via its natural transduction route, which would involve clathrin-mediated endocytosis, endosomal escape, and nuclear translocation through the nuclear pore complex. Instead, the amplicon is introduced into the cell using cationic polymers, leaving only a minor fraction to reach the nucleus, the place where AdV genome replication occurs. Second, naturally transduced AdV genomes are condensed with AdV proteins (e.g., the histone-like protein VII) and they carry a covalently bound terminal protein (TP) at both 5’-ends, which protects the genome from exonucleases and facilitates correct genome localization and potent induction of genome replication.

To address these technical limitations and study the components of the AdV DNA replication machinery in a more natural context, we developed an improved *trans*-replication assay. To this end, the host cells were first transfected with a single vDNA or a combination of plasmids intended to be probed for reconstituting a functional AdV replication machinery. Subsequently, the cells were transduced with replication-deficient HC-AdV-C5 virions, encoding only a GFP reporter gene. Successful assembly of the AdV replication machinery by the transfected DNA should result in *trans*-amplification of the incoming, transduced HC-AdV genome (Figure 3A). To validate this refined assay system, HEK293 cells were pre-transfected with non-coding pC4HSU vDNA, followed by transduction with HC-AdV particles encoding only a GFP reporter cassette. Following qPCR analysis of total DNA extract, there were no differences in HC-AdV genome copy numbers detectable at 4 h and 48 h post-transduction (Figure S5), highlighting that HC-AdVs lack inherent genome replication. In contrast, pre-transfection with ΔE1,ΔE3 vDNA confirmed that the encoded replication machinery promotes HC-AdV genome *trans*-amplification, also when the viral chromosome is delivered via the natural AdV transduction. Normalization as x-fold amplification per cell underscored that ΔE1,ΔE3 vDNA drives more than 300-fold amplification of the transduced HC-AdV genome per cell (Figure 3B). Unlike in the amplicon-based *trans*-replication system, additional deletion of the E4 TU did not fully abolish replication, as pre-transfection with ΔE1,ΔE3,ΔE4 displayed only reduced replication efficiency in comparison ΔE1,ΔE3 vDNA. Yet, additional expression of E4 ORF6 rescued the AdV replication to its maximal capacity again, as defined by pre-transfection with the positive control ΔE1,ΔE3 vDNA. Moreover, the combined expression of the three E2 proteins (DBP, pTP, Ad Pol) with E4 ORF6 was sufficient to drive more than 2,000-fold replication of transduced HC-AdV genomes in HEK293 cells.

**Figure 3.**
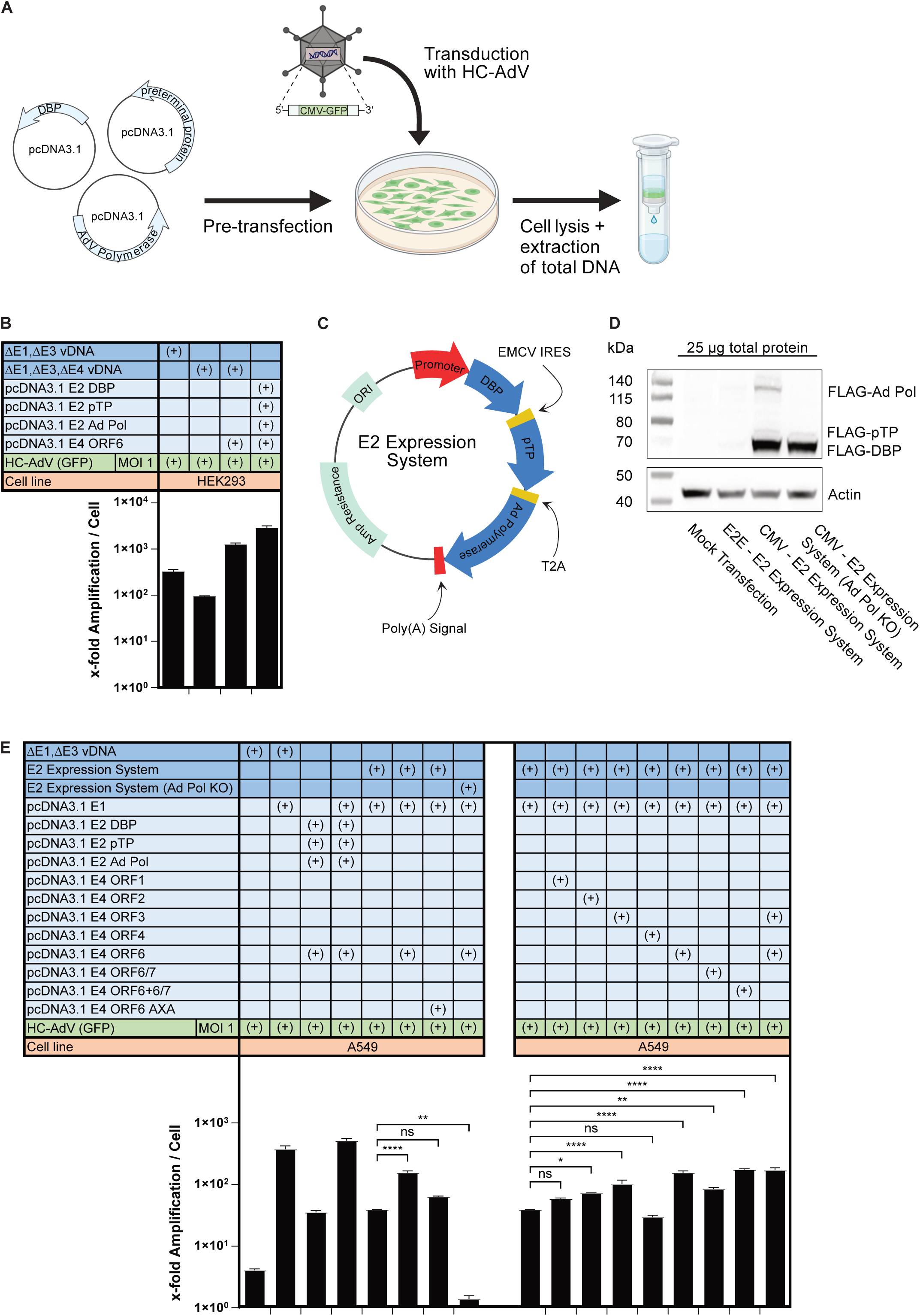
Designed E2 expression system is functional and together with E4 ORF6 reconstitutes a minimal replication system promoting amplification of incoming, transduced HC-AdV genomes. (A) Graphical depiction of a *trans*-replication assay to quantify amplification of incoming, transduced HC-AdV genomes. Host cells were pre-transfected with either an individual or a combination of expression plasmids (light blue) that are being probed for reconstitution of a functional AdV DNA replication machinery. Pre-transfected cells were transduced with HC-AdV (GFP) (gray), and changes in HC-AdV genome copy numbers were quantified 48 h post-transduction via qPCR upon extraction of total DNA. (B) qPCR quantification of x-fold replication of incoming, transduced HC-AdV genomes upon pre-transfection of HEK293 cells with AdV replication machineries (dark blue) or AdV genes (light blue). *Trans*-replicated HC-AdV genomes per cell were determined and normalized to non-replicated HC-AdV genomes per cell 48 h post-transduction. (C) Schematic illustration of designed E2 expression system, comprising a tricistronic expression cassette that supplies all three E2 proteins (DBP, pTP, Ad Pol; depicted in dark blue) in FLAG-tagged form. Promoter and polyadenylation signal are shown in red and translation-mediating motifs in yellow. (D) Western blot analysis and detection of E2 proteins by FLAG-tag immunostaining confirmed functional expression of E2 proteins 36 h post-transfection of HEK293 cells with different E2 expression systems. β-actin was immunostained as a loading control. (E) Quantification of x-fold amplification of incoming, transduced HC-AdV genomes via qPCR, as analyzed 48 h post-transfection of A549 cells with different replication systems (ΔE1,ΔE3 vDNA, E2 expression systems; shown in dark blue) or a combination of individual AdV expression plasmids encoding E1, E2 and E4 genes (light blue). Statistics: Representative data of two to three independent experiments are shown. Bar graphs represent mean of x-fold amplification ± SD, n = 3. Statistical significance was determined by one-way ANOVA with Dunnett’s test for multiple comparisons to the control group of co-transfected E2 expression system and E1. Not significant (ns) P > 0.05; *P ≤ 0.05; **P ≤ 0.01; ***P ≤ 0.001; ****P ≤ 0.0001.

Developing a genome-replicating HC-AdV requires encoding all necessary AdV genes in *cis*, but we also prioritized maximizing the transgene capacity to allow increased *in situ* expression of multiple therapeutic proteins. In light of the complex genetic architecture of the WT AdV-C5 (Figure S1), fundamental rearrangements of the involved AdV genes were required to generate a robust, compact, and splicing-independent minimal replication system. A tricistronic E2 expression system was designed which transcribes all three E2 proteins under control of either the endogenous E2 early (E2E) or a constitutive CMV promoter (Figure 3C). As this forms one single transcript with DBP upstream as the quantitatively most needed component, cap-independent translation of pTP was enabled using an internal ribosomal entry site (IRES), while Ad Pol is translated using a T2A self-cleavage (ribosome skipping) peptide. To monitor functionality of this system we encoded all E2 proteins as FLAG-tagged variants. As a control, we additionally generated a replication-deficient, Ad Pol knock-out (Ad Pol KO) version of the E2 expression system, lacking the T2A-Ad Pol sequence.

We next wanted to validate expression of FLAG-tagged E2 proteins and therefore transfected HEK293 cells with the different E2 expression systems. Western blot analysis and immunostaining demonstrated that the E2E promoter drives only very weak E2 expression while the CMV promoter results in potent expression of DBP, pTP and Ad Pol (Figure 3D). Notably, DBP is expressed much more abundantly compared to pTP and Ad Pol, and thus the CMV – E2 expression system is closely mimicking the stoichiometry of the three E2 proteins observed upon natural AdV transduction.^81^ Deletion of the Ad Pol sequence from the E2 expression system (CMV – E2 expression system (Ad Pol KO)) resulted in the absence of detectable FLAG-Ad Pol expression.

Next, we analyzed whether the developed CMV – E2 expression system promotes functional replication of incoming, transduced HC-AdV genomes in A549 cells using our *trans*-replication assay. HC-AdV genome quantification via qPCR confirmed that sole transfection of ΔE1,ΔE3 vDNA failed to initiate replication, however, after co-transfection with E1 robust amplification was observed (Figure 3E). DBP, pTP, Ad Pol and E4 ORF6 alone induced some HC-AdV genome amplification, though maximum replication was only achieved in the presence of E1. Importantly, our developed CMV – E2 expression system induced more than 150-fold HC-AdV genome replication when combined with E4 ORF6 and E1, being only slightly less efficient in comparison to the positive control ΔE1,ΔE3 vDNA + E1.

To investigate if E4 ORF6 enhances replication through association with E1B-55k or via an alternative mechanism, we co-transfected A549 cells with CMV – E2 expression system, E1 and a E4 ORF6 mutant termed E4 ORF6 AXA.^82^ This mutant features two mutations (R243A and L254A), disrupting the putative RXL motif and thus preventing it from interacting with E1B-55k, while retaining its ability to interact with p53.^82^ Co-transfection with E4 ORF6 AXA resulted in reduced replication efficiency, suggesting that the interaction between E1B-55k and E4 ORF6 is indeed required for full AdV genome replication efficiency. Transfection with the E2 expression system lacking Ad Pol (Ad Pol KO) failed to support replication, confirming that Ad Pol, and not cellular polymerases, is required for replication. Screening of different E4 gene products in combination with the CMV – E2 expression system and E1 validated our previous finding that the most potent replication is driven by E1, E2, and E4 ORF6. Additional *trans*-complementation with E4 ORF3 or with relevant intermediate and late genes did not further enhance HC-AdV genome replication (Figure S6), corroborating our earlier observations that intermediate and late gene products are not required for AdV replication.

We thus identified in total seven AdV proteins critical for promoting the replication of incoming, transduced HC-AdV genomes in E1 non-complementing cells. Additionally, we have designed a compact and splicing-independent E2 expression system that ensures robust E2 protein production, resulting in replication levels nearly comparable to the positive control ΔE1,ΔE3 vDNA + E1.

### HC-AdV *trans*-replication results in increased payload expression of a vector-encoded fluorescent reporter

To assess whether HC-AdV genome replication indeed leads to increased expression of the vector-encoded payload, we first performed similar *trans*-replication assays in HEK293T cells and then additionally quantified reporter expression levels of the HC-AdV-encoded GFP reporter via flow cytometry by measuring the mean fluorescence intensity (MFI). HEK293T cells were pre-transfected with non-coding vDNA followed by transduction with HC-AdV (GFP) virions at an MOI of 1. GFP was evenly expressed 48 h post-transduction and no signs of a cytopathic effect were observable, as indicated by brightfield and fluorescent microscopy (Figure 4A). The MFI could be determined to 1376, which represents the baseline GFP expression levels without any genome replication. Pre-transfection of HEK293T cells with ΔE1,ΔE3 vDNA induced replication of the HC-AdV (GFP) genome, resulting in enhanced GFP expression, as demonstrated by fluorescence microscopy and a ∼2.4-fold increased MFI. As ΔE1,ΔE3 vDNA supplies relevant AdV proteins, induction of cytopathic effect was noticeable in brightfield microscopy. Further deletion of the E4 TU (ΔE1,ΔE3,ΔE4 vDNA) reversed both effects, and no cytotoxicity but also no significantly increased MFI was detectable any longer. When ΔE1,ΔE3,ΔE4 vDNA was *trans*-complemented with E4 ORF6, GFP expression increased markedly, as shown by fluorescent microscopy, and as reflected by a ∼3.4-fold increased MFI and a pronounced shift in the GFP^+^ population upon flow cytometric analysis. Complementing E4 ORF6 expression moreover restored induction of the cytopathic effect. Co-transfection of HEK293T cells with only four AdV genes (DBP, pTP, Ad Pol, E4 ORF6) was sufficient to significantly increase GFP expression as illustrated by fluorescence microscopy and the distinct shift in the GFP^+^ population resulting in a ∼2.2-fold increase in MFI. Notably, there was no sign of induction of an observable cytopathic effect, suggesting that the individual components of the minimal replication system are indeed not toxic to the host cell.

**Figure 4.**
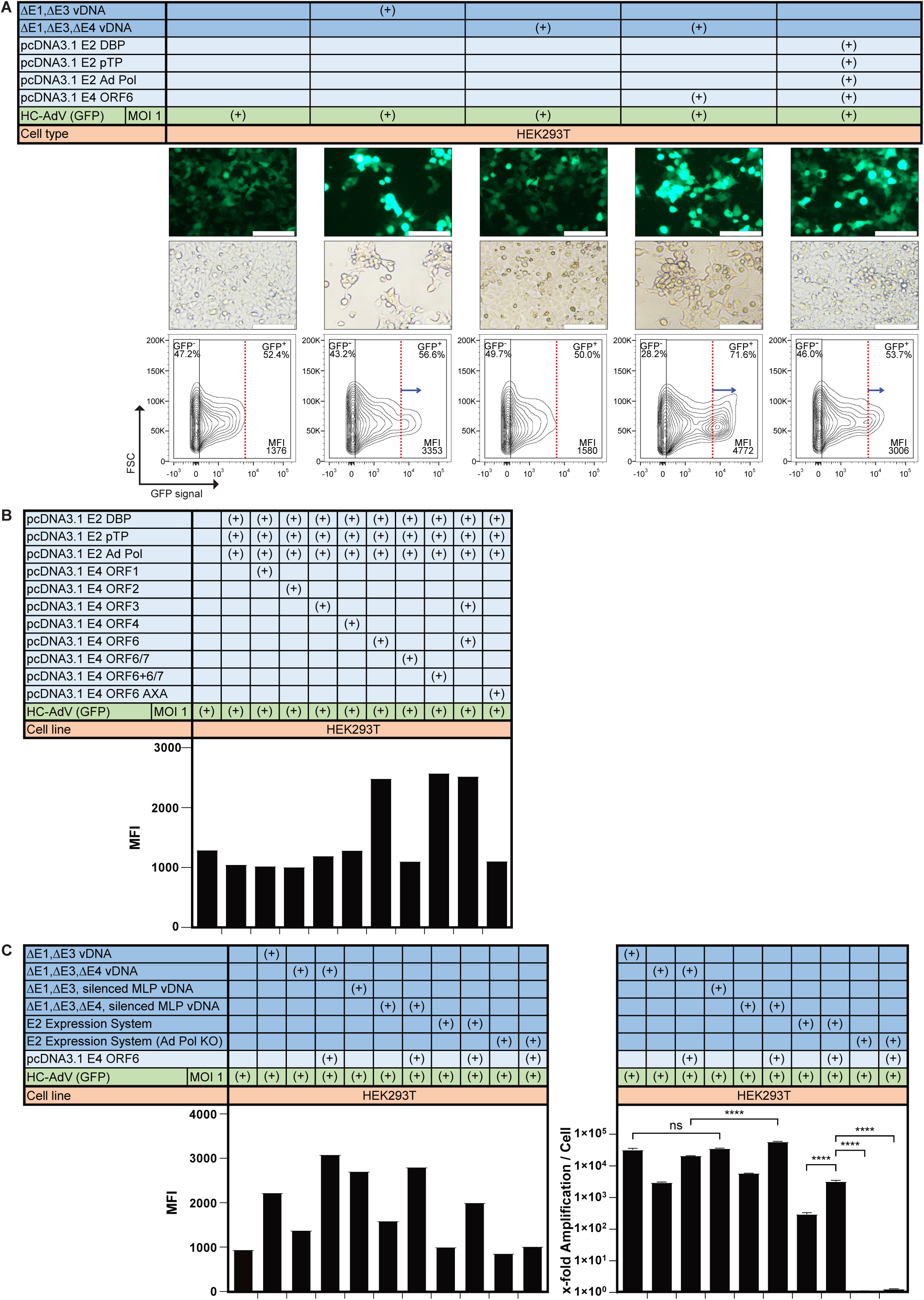
*Trans*-replication of incoming, transduced HC-AdV genomes promotes increased expression of the vector-encoded reporter in HEK293T cells. (A) Representative bright-field and fluorescent micrographs of HEK293T cells, pre-transfected with different AdV replication machineries (dark blue) or AdV genes (light blue), analyzed 72 h post-transduction with HC-AdV (GFP) at an MOI of 1. Fluorescent reporter expression was quantified via flow cytometric analysis and displayed as mean fluorescent intensity (MFI). Enhanced GFP expression following replication was evident as a shift in the GFP^+^ population, as highlighted by the arrow (blue) in the contour plots. Scale bars = 100 µm. (B) Flow cytometric analysis of GFP reporter expression levels after co-transfection of HEK293T cells with expression plasmids encoding DBP, pTP, or Ad Pol in combination with different genes of the E4 TU and transduction with HC-AdV (GFP) (MOI 1). Reporter expression levels were analyzed 72 h post-transduction and quantified as MFI. (C) Quantification of reporter expression by flow cytometry (left panel) after pre-transfection of HEK293T cells with different AdV replication systems (dark blue) and an expression plasmid encoding E4 ORF6 (light blue), followed by transduction with HC-AdV (GFP) virions (MOI 1). Cells were pre-transfected with ΔE1,ΔE3 vDNA or ΔE1,ΔE3,ΔE4 vDNA (with each of the vDNAs additionally encoding a silenced major late promoter (MLP)) in combination with E4 ORF6, and fluorescence was determined 72 h post-transduction (left panel) and genome replication was additionally quantified via qPCR and displayed as x-fold amplification per cell (right panel). Statistics: Representative data of two independent experiments are shown. Bar graphs represent mean of x-fold amplification ± SD, n = 3, or absolute MFI. Statistical significance was determined by one-way ANOVA with Dunnett’s test for multiple comparisons. Not significant (ns) P > 0.05; *P ≤ 0.05; **P ≤ 0.01; ***P ≤ 0.001; ****P ≤ 0.0001.

While E4 ORF6, in combination with DBP, pTP, Ad Pol, and all three E1 proteins (provided by the HEK293T host cell) drives the most potent HC-AdV genome replication we wondered if other E4 TU gene products, besides E4 ORF6, also enhance payload expression. Therefore, we *trans*-complemented DBP, pTP, and Ad Pol with different E4 TU gene products and transduced the HEK293T cells with HC-AdV (GFP) virions. Flow cytometric analysis demonstrated that only E4 ORF6 is able to significantly increase payload expression (∼2-fold), while other E4 gene products (ORF1, 2, 3, 4, and 6/7) do not mimic this effect (Figure 4B). As transfection with an expression plasmid encoding E4 ORF6+6/7 supplies both E4 ORF6 and the fusion protein E4 ORF6/7, the phenotype of increased GFP expression was most likely attributed to the presence of E4 ORF6. Similarly to AdV genome replication, E4 ORF6’s ability in enhancing payload expression is strongly dependent on its interaction with E1B-55k. Upon *trans*-complementation with the E4 ORF6 AXA mutant, the robust increase in GFP expression was abolished and the MFI was comparable again to that observed in the absence of HC-AdV genome replication.

Although late genes are not required for AdV genome replication, we wanted to investigate potential multifunctional roles and assess whether they might contribute to enhanced transgene expression. Therefore, we pre-transfected HEK293T cells with ΔE1,ΔE3 vDNA and ΔE1,ΔE3,ΔE4 vDNA, both as versions encoding either an active or silenced MLP. In line with our previous data from *trans*-replication assays, transfection with ΔE1,ΔE3 vDNA resulted in ∼2-fold increase in payload expression, while deletion of the E4 TU drastically reduced this effect (Figure 4C, left panel). As seen before, *trans*-complementation with E4 ORF6 restored enhanced payload expression (∼3-fold increase in MFI). Similar observations were made when the MLP was silenced on these vDNAs, indicating that late gene expression (driven by the MLP) is in fact not necessary for an increased payload expression upon HC-AdV genome replication. Additionally, when HEK293T cells were pre-transfected with our E2 expression system alongside E4 ORF6, we observed a ∼2-fold increase in MFI upon transduction with HC-AdV (GFP) virions. This suggests that, despite the constitutive CMV promoter and the tricistronic expression cassette, our E2 expression system phenotypically resembles the activity of the E2 genes when present in their native configuration on ΔE1,ΔE3 vDNA. As expected, t*rans*-complementation with the replication-deficient E2 expression system (Ad Pol KO) did not increase the MFI.

Simultaneous qPCR analysis highlighted that ΔE1,ΔE3 vDNA (active and silenced MLP) promoted over ∼30,000-fold amplification in HEK293T cells (Figure 4C, right panel). Deletion of the E4 TU consequently reduced replication efficiency which was rescued for both versions, encoding the active and inactive MLP, to similar levels compared to ΔE1,ΔE3 vDNA upon *trans*-complementation with E4 ORF6. Replication of HC-AdV genomes mediated by the E2 expression system was highly dependent on E4 ORF6, with replication increasing from ∼300-fold to ∼3,000-fold upon co-expression of E4 ORF6. The Ad Pol-free E2 expression system (Ad Pol KO) did not induce HC-AdV replication, regardless of the presence of E4 ORF6. Of note, transfection of ΔE1,ΔE3,ΔE4 vDNA and of the E2 expression system in conjunction with E4 ORF6 resulted in comparable levels of HC-AdV genome replication (Figure 4C, right panel), however only the latter mediated a significantly increased payload expression (Figure 4C, left panel), suggesting that both replication and E4 ORF6 activity is required to enhance transgene expression.

In conclusion, using our *trans*-complementation assay, we demonstrated that AdV genome replication leads to increased transgene expression. In HEK293T cells, only four AdV gene products – DBP, pTP, Ad Pol, and E4 ORF6 – are essential for replication and to thus enhance payload expression. We identified two key prerequisites for this effect, (i) effective AdV genome replication must occur, and (ii) the presence of E4 ORF6, and particularly its interaction with E1B-55k. Additionally, we confirmed that our E2 expression system efficiently drives HC-AdV genome replication and boosts payload expression when combined with E4 ORF6. Our designed minimal replication system thus comprises three transcriptional units and in total seven genes: the WT E1 TU (expressing E1A, E1B-19k, and E1B-55k), our E2 expression system (supplying DBP, pTP, and Ad Pol), and an E4 ORF6 expression cassette.

### HC-AdV encoding a minimal replication system results in *cis*-replication and increased reporter expression upon transduction of E1-non-complementing BT-474 cells

To test whether the developed minimal replication system promotes both *cis*-acting amplification and enhanced payload expression, we cloned and produced three different HC-AdVs (Figure 5A). This included two control vectors that encode (i) only a GFP reporter cassette (HC-AdV (GFP)) or (ii) a GFP reporter cassette and additionally the native E1-coding region of the AdV-C5 (HC-AdV (E1 + GFP)). The latter was designed to discern whether elevated payload expression indeed results from genome replication, or is rather a consequence of promiscuous E1-mediated host transcription factor activation. The third vector, HC-AdV (GFP + Minimal Replication System), encoded a GFP reporter and the optimized, final minimal replication system, comprising E1, an E2 expression system, and the E4 ORF6 expression cassette. To enhance replication, the E2 expression system had been further optimized and DBP was now expressed separately from a CMV promoter, while pTP and Ad Pol were transcribed as one dicistronic ORF using a PGK promoter and T2A self-cleavage peptide (Figure S7A, “Minimal Replication System 3”). Elevated DBP levels were shown to increase genome replication (Figure S7B and S7C). The endogenous E4 promoter provided sufficient activity to express levels of E4 ORF6 required for replication (Figure S8).

**Figure 5.**
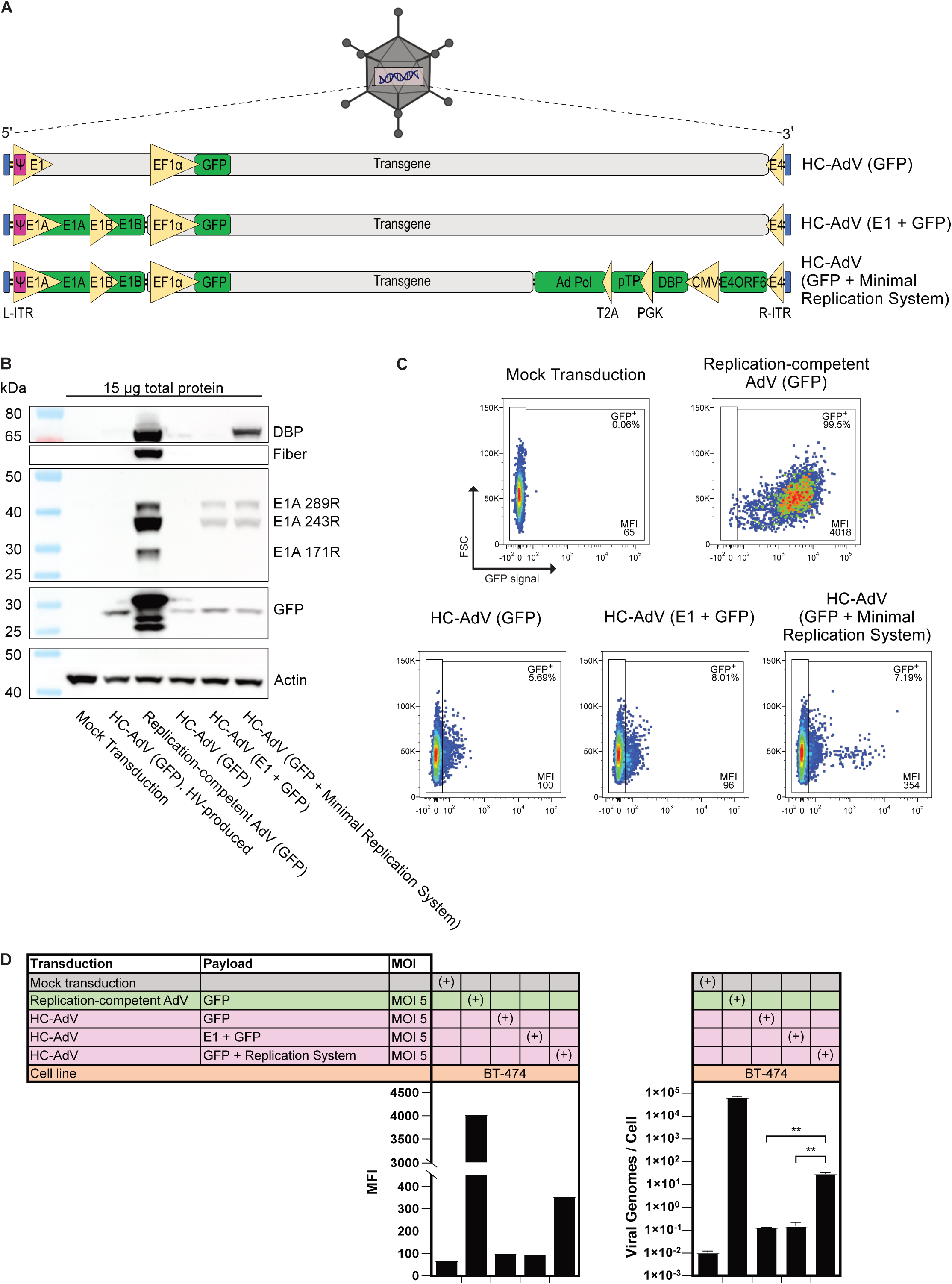
Transduction of BT-474 cells with genome-replicating HC-AdV results in increased reporter expression levels. (A) Schematic depiction of three HC-AdV vectors that were designed to assess functionality of genome-replicating HC-AdV vectors. Promoters and T2A self-cleavage peptide are depicted in yellow and protein coding regions are shown in green. Transduction with HC-AdV (GFP) provides benchmark expression of GFP without genome replication, and HC-AdV (E1 + GFP) allows evaluating any potential increase in reporter expression due to E1-mediated *trans*-activation of cellular transcription factors. Genome-replicating HC-AdV (GFP + Minimal Replication System) encodes a functional replication machinery comprising expression cassettes for E1, E2 and E4 ORF6. All three HC-AdVs were produced using helper plasmids to exclude HV contamination. (B) Western blot analysis and immunostaining of selected proteins 36 h after transduction of E1-non-complementing BT-474 cells with HV-produced HC-AdV (GFP), RC-AdV (GFP), or helper plasmid-produced HC-AdVs (GFP), (E1 + GFP), or (GFP + Minimal Replication System) (MOI 5). Detection of β-actin served as a loading control. (C) Flow cytometric analysis of GFP^+^ populations 72 h post-transduction (MOI of 5) of BT-474 cells with RC-AdV (GFP) or HC-AdVs (GFP), (E1 + GFP), or (GFP + Minimal Replication System). (D) Quantification of reporter expression levels, displayed as mean fluorescent intensity (MFI) after flow cytometric analysis, and viral genomes per cells (qPCR analysis) 72 h post-transduction (MOI of 5) of BT-474 cells with either RC-AdV (GFP) (green) or different HC-AdV vectors (purple). Statistics: Representative data of two independent experiments are shown. Bar graphs represent mean AdV genome copy numbers ± SD, n = 3, or absolute MFI. Statistical significance was determined by one-way ANOVA with Dunnett’s test for multiple comparisons. Not significant (ns) P > 0.05; *P ≤ 0.05; **P ≤ 0.01; ***P ≤ 0.001; ****P ≤ 0.0001.

Standard HC-AdV vector production workflows employ HVs (modified ΔE1,ΔE3 first-generation AdV vectors), which are rendered non-packable in the producer cell. However, this process is imperfect, leading to <1% HV contamination per HC-AdV preparation, and occasional co-transduction of the same cell with both HC-AdV and HV. Since HVs are typically E1-deleted and non-replicative, contamination poses no problem, unless the HC-AdV vector encodes a functional E1 TU. Then, it can *trans*-activate the life cycle of the co-transduced HV, resulting in HV progeny formation. Given that our HC-AdV (GFP + Minimal Replication System) encodes E1, we thus generated all three HC-AdV vectors described above by using helper plasmids instead of HV during production.^83^ This ensured that no contaminating HV is co-produced during generation of HC-AdVs and we thus excluded potential *trans*-activation of HV by E1-encoding HC-AdVs during subsequent testing of our produced vectors.

To confirm the genome identity of our produced HC-AdV vectors, we first transduced E1-non-complementing BT-474 cells at an MOI of 5, alongside RC-AdV (GFP), and analyzed protein expression 48 h post-transduction via western blotting and immunostaining. BT-474 cells exhibit a slow doubling time of ∼80 h^84^ and therefore represent an ideal cell system to characterize AdV genome replication as cell division does not interfere with genome amplification. For comparison, BT-474 cells were additionally transduced with conventionally, HV-produced HC-AdV (GFP), which is identical to helper plasmid-produced HC-AdV (GFP) virions. Since the HV-produced HC-AdV (GFP) is devoid of all AdV genes, it only expresses the transgene GFP (Figure 5B). Transduction of BT-474 cells with RC-AdV (GFP) resulted in potent expression of GFP, along with three detectable E1A isoforms, DBP, and fiber, confirming productive infection. As expected, HC-AdV (GFP) produced with helper plasmid (and not HV) induced identical protein expression patterns as monitored after transduction with HV-produced HC-AdV (GFP), thus validating our helper plasmid-based HC-AdV production system. While western blot analysis after transduction of BT-474 cells with HC-AdV (E1 + GFP) or HC-AdV (GFP + Minimal Replication System) demonstrated expression of GFP, E1A, or GFP, E1A, and DBP, respectively, we identified reduced E1A levels compared to RC-AdV transduction, with the E1A isoform 171R absent. Fiber expression was undetectable after transduction with all HC-AdVs, confirming that undesired formation of RC-AdVs via homologous recombination did not occur during production.

We then transduced BT-474 cells (MOI of 5), harvested the cells after 72 h, and analyzed payload expression via flow cytometry and genome replication using qPCR. Transduction with RC-AdV (GFP) resulted in nearly all cells being GFP^+^ while displaying pronounced GFP expression (MFI of ∼4000), demonstrating a strikingly enhanced payload expression due to AdV genome replication (Figure 5C and 5D, left panel). Transduction with HC-AdV (GFP) or HC-AdV (E1 + GFP) resulted in weak GFP expression, with MFIs of 100 and 96, respectively. In contrast, transduction of BT-474 cells with HC-AdV (GFP + Minimal Replication System) resulted in a ∼3.5-fold increase in MFI, indicating significantly enhanced GFP expression in transduced cells. In comparison to non-genome-replicating HC-AdVs, such as (GFP) or (E1 + GFP), transduction with our genome-replicating HC-AdV generated a distinct population of high GFP-expressing cells (Figure 5C), confirming the concept and functionality of our developed genome-replicating HC-AdV.

Independent AdV genome copy number quantification by qPCR analysis confirmed more than ∼100,000-fold genome replication upon transduction of BT-474 cells with RC-AdV (GFP) (Figure 5D, right panel). Transduction with HC-AdV (GFP) and (E1 + GFP) resulted in ∼0.1 vector copies/cell 72 h post-transduction, while transduction with our genome-replicating HC-AdV (GFP + Minimal Replication System) displayed a vector copy number of ∼30 genomes/cell at end point. This suggests that our minimal replication system is indeed functional and promotes more than 300-fold *cis*-acting replication of delivered genomes per cell.

Although the minimal replication system in this proof-of-concept study did not fully match the RC-AdV (GFP) in genome replication and payload expression, it nonetheless promoted ∼300-fold genome replication and enhanced payload expression, as demonstrated by a high GFP-expressing population. In summary, our presented data are indicative of the functionality of our minimal replication system, as it promotes *cis*-replication and drives a 3.5-fold increase in MFI. Additional expression of E1 (HC-AdV (E1 + GFP)) did not measurably affect payload expression levels in BT-474 cells.

### HC-AdV encoding the minimal replication system strongly boosts payload expression in E1-complementing HEK293T, with no further increase by *trans*-complementation

We next performed similar transduction assays using E1-complementing HEK293T as host cells and analyzed GFP expression levels and AdV copy numbers 72 h post-transduction. While transduction with RC-AdV (GFP) resulted in strong GFP expression (MFI ∼5000) (Figure 6A, left panel), HC-AdV (GFP) and HC-AdV (E1 + GFP) induced only low reporter expression (MFI of ∼130 and ∼100, respectively). In comparison, HC-AdV (GFP + Minimal Replication System) drastically increased GFP levels by ∼20-fold, as reflected by an elevated MFI of ∼2450. Quantification via qPCR indicates robust genome replication for RC-AdV (∼100,000 copies/cell), while transduction with both control vectors, HC-AdV (GFP) and (E1 + GFP) resulted in very few genomes in the host cell (∼0.06 and ∼0.02 copies/cell, respectively) at end point. Upon transduction with HC-AdV (GFP + Minimal Replication System) we detected ∼160 copies/cell, indicating potent (∼4,000-fold) *cis*-acting HC-AdV genome replication.

**Figure 6.**
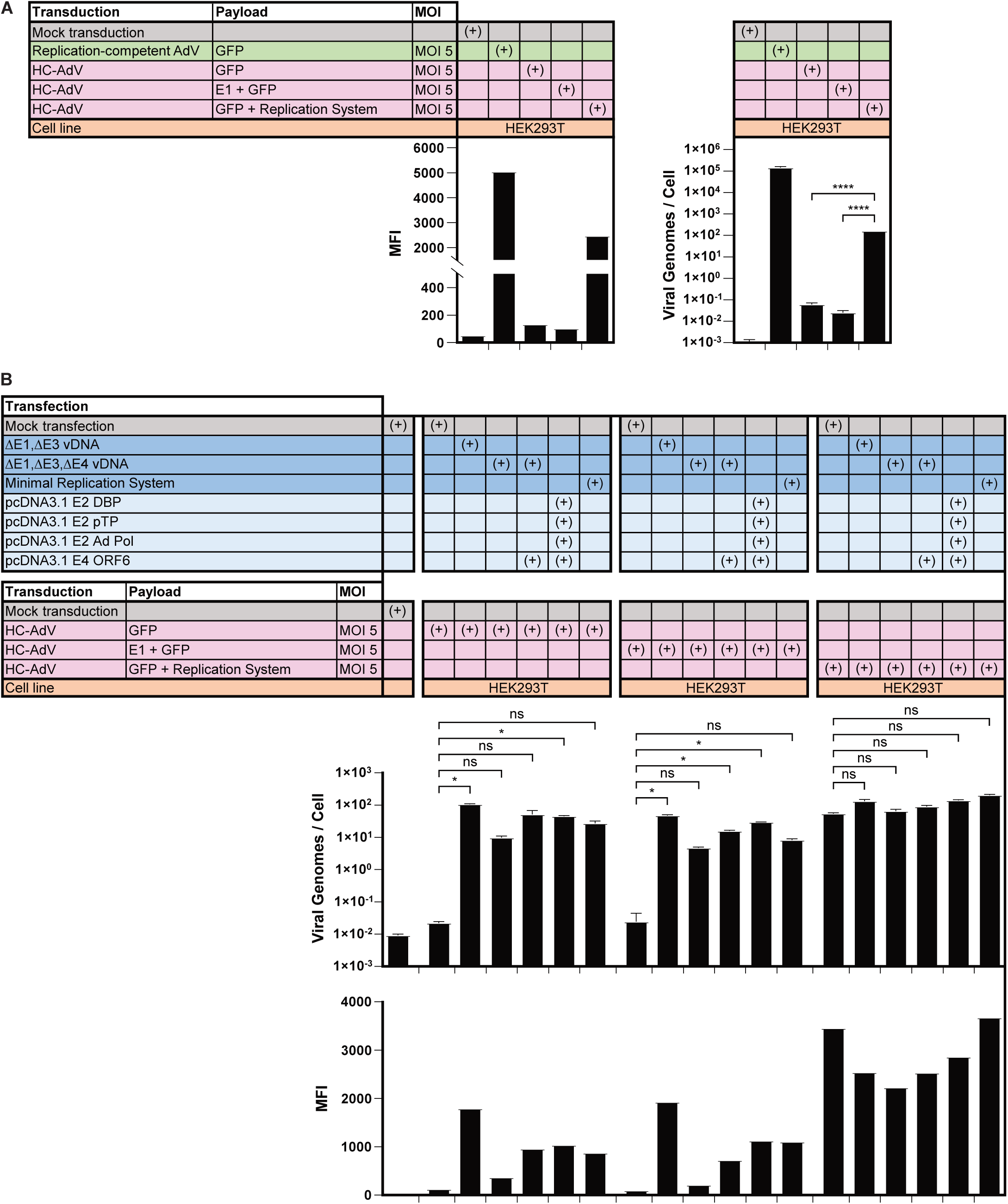
Transduction of HEK293T cells with genome-replicating HC-AdV reconstitutes a fully functional minimal replication system with no further increase in activity by *trans*-complementation. (A) Quantification of reporter expression levels, displayed as mean fluorescent intensity (MFI) after flow cytometric analysis, and viral genomes per cells (qPCR analysis) 72 h post-transduction (MOI of 5) of E1-complementing HEK293T cells with RC-AdV (GFP) (green) or different HC-AdV vectors (purple). (B) AdV vector copy quantification by qPCR and flow cytometric analysis of GFP reporter expression 72 h post-transduction (MOI of 5) of HEK293T cells with HC-AdV vectors (purple). Cells were pre-transfected 4 h prior to transduction with indicated AdV replication system (dark blue) or a combination of individual AdV expression plasmids encoding E2 and E4 ORF6 genes (light blue). Representative data of two independent experiments are shown. Bar graphs represent mean AdV genome copy numbers ± SD, n = 3, or absolute MFI. Statistical significance was determined by one-way ANOVA with Dunnett’s test for multiple comparisons. Not significant (ns) P > 0.05; *P ≤ 0.05; **P ≤ 0.01; ***P ≤ 0.001; ****P ≤ 0.0001.

To assess if expression and assembly of the encoded minimal replication system is fully functional after HC-AdV genome delivery, we next performed *trans*-complementation assays in HEK293T cells. For this purpose, we first transfected the cells with vDNAs or different expression plasmids encoding individual AdV genes, followed by transduction with our HC-AdV vectors. As previously observed, mock-transfected HEK293T cells transduced with either HC-AdV (GFP) or (E1 + GFP) displayed only very few vector genomes/cell (∼0.2 copies/cell) (Figure 6B, right panel) and low GFP expression (MFI ∼105). Pre-transfection with different, functional AdV replication systems established potent genome amplification and up to ∼18-fold increased MFI. Transduction with HC-AdV (GFP + Minimal Replication System) upon mock transfection resulted in ∼2,300-fold replication and elevated GFP levels, indicating that genome amplification was entirely promoted by the *cis*-encoded replication system. Importantly, no further increase in genome replication or GFP expression was observed upon pre-transfection of AdV replication machineries, suggesting that the *cis*-encoded system supplies sufficient protein levels of the replication machinery to function at its maximum capacity.

Altogether, these data demonstrate that genome-replicating HC-AdV vectors can be an efficient approach to enhance payload expression. By encoding only seven AdV genes, we showed that these genes are sufficient to assemble a functional replication machinery, leading to vector genome amplification in the nucleus of the host cell and increased *in situ* payload expression. Additional *trans*-complementation did not further enhance replication or transgene expression, indicating that the minimal replication system induces maximal genome replication. Importantly, this compact replication system retains over 22 kb of transgene capacity on HC-AdV, providing ample space to deliver therapeutic proteins and enabling the expression of multiple biologics for combination therapy.

## Discussion

Conventional HC-AdV vectors lack all AdV genes, providing a large transgene capacity, reduced cytotoxicity, and prolonged transgene expression due to the absence of AdV protein expression. However, the lack of genome replication limits *in situ* payload expression, which is higher with RC-AdV or SC-AdV vectors. This proof-of-concept study presents a new, genome-replicating HC-AdV vector class that combines the benefits of both HC-AdV and RC-AdV while minimizing their shortcomings. We developed two transfection-based *trans*-replication cell assays, enabling us to systematically screen AdV genes for their contribution towards genome replication and to remove all replication-unrelated genes. By encoding only seven AdV genes on an HC-AdV vector, our vector promotes potent vector genome amplification in *cis*, and thus generates a larger pool of DNA templates for increased transcription and translation of the encoded payload. Deletion of AdV proteins reduced cytotoxicity and provided a large transgene capacity of more than 22 kb, allowing increased *in situ* expression of combinatorial protein therapeutics for complex indications such as cancer.

For more than four decades it has been known that *in vitro* AdV genome replication requires only the three AdV E2 proteins that act in conjunction with three cellular factors.^52,53^ However, the contribution of other AdV gene products, e.g., from the E1 and E4 TUs, towards efficient replication in cells, and particularly their roles in modulating the host cell environment to support potent AdV genome replication *in vivo*, remained partially elusive. We therefore screened 24 out of the more than 40 described AdV-C5 genes, allowing us to identify a combination of seven AdV gene products specifically required for AdV genome replication and enhanced payload expression in E1-non-complementing cells. These include the gene products from the E1 TU (E1A, E1B-19k, and E1B-55k), all three gene products from the E2 TU (DBP, pTP, Ad Pol), and E4 ORF6 (Figure 2B + 3E).

Previous studies with oncolytic AdV (OAdV) vectors displaying deletions of E1B-55k suggested that E1B-55k is not per se required for AdV genome replication, especially when targeting cancer cells.^85^ Given the proliferative nature of cancer cells and that expression of our minimal replication does not rely on E1A-dependent promoters, we initially anticipated overcoming the necessity of encoding the entire E1 TU on our genome-replicating HC-AdV, particularly in consideration of preventing the transformation potential of E1A. Nevertheless, both of our cell-based *trans*-replication assays clearly indicate that the E1 proteins are essential for AdV vector genome replication (Figure 2B + 3E). Due to a mutation in the CDKN2B gene (also called p15^INK4^), A549 cells display a defective Rb pathway^86–88^ and are therefore not dependent on E1A activity for cell cycle progression. This suggests that at least one or multiple alternative functions of E1A are required for AdV genome replication, such as modulation of a diverse set of host cell transcription factors or chromatin reorganization.^78^ While we did not further investigate the roles of the individual E1 proteins (E1A, E1B-19k, and E1B-55k) or E1A isoforms, we presume that E1B-19k and 55k are most likely essential when E1A is present to counteract the apoptotic stimuli induced by E1A and AdV genome replication.^78^

We demonstrated that the E4 gene products ORF3 and ORF6 support replication of incoming, transduced HC-AdV genomes, with E4 ORF6 being most effective when combined with E1A, E1B, DBP, pTP, and Ad Pol (Figure 3E). Consistent with earlier studies,^60,61^ either E4 ORF3 or ORF6 are necessary for rescuing genome replication and progeny formation upon ΔE4 AdV-C5 transduction, primarily through suppression of the host cell’s DNA damage repair (DDR). Specifically, the MRN complex (MRE11-Rad50-NBS1) and DNA ligase IV recognize the AdV genome as damaged DNA, promoting end-to-end ligation of viral DNA into concatemers, perturbing viral DNA replication and packaging into progeny particles.^80^ E4 ORF3 and ORF6 exhibit functional redundancy but employ different DDR inhibition mechanisms: E4 ORF3 sequesters DDR components in PML-NBs and thus spatially separates them from AdV replication compartments (RCs),^56^ while E4 ORF6 drives proteasomal degradation of host factors via formation of an E3 ubiquitin ligase complex.^67^ RCs are virus-induced, membrane-less, nuclear sub-compartments, likely formed through liquid-liquid phase separation. These structures concentrate essential viral and host proteins, promoting AdV gene transcription, DNA replication, and progeny formation while excluding inhibiting factors.^56,89,90^

The distinct mechanisms of DDR inhibition employed by E4 ORF3 and ORF6 may explain why both individually support replication of incoming, transduced HC-AdV genomes (Figure 3E), whereas only E4 ORF6 promotes replication of transfected AdV mini-chromosomes (Figure 1D). Notably, incoming, transduced HC-AdV genomes are covalently attached to two copies of terminal protein (TP) and condensed with protein VII, whereas the bacterially produced AdV mini-chromosome is protein-free. TP is crucial for mediating the nuclear localization and matrix attachment of incoming HC-AdV genomes, and robust initiation of viral replication.^91–93^ E4 ORF3 polymerization redistributes nuclear matrix-associated PML-NBs^94^ near AdV RCs and thus physically separates RCs from components of the DDR.^56,95^ It is thus conceivable that TP-mediated nuclear matrix attachment of the AdV genome is necessary for E4 ORF3 to effectively suppress DDR responses and hence enables efficient AdV genome replication. In contrast, E4 ORF6 inhibits DDR independently of the nuclear matrix and can therefore promote replication of both, transduced and transfected AdV genomes.

Upon studying the role of the E4 proteins in increasing payload expression following HC-AdV genome replication, we observed that genome amplification alone does not fully account for, but rather is a prerequisite for elevating expression levels of the encoded transgene. The effects of E4 ORF3 and ORF6 on supporting enhanced expression were found to be very different. Although E4 ORF3 also supports AdV genome replication — albeit to a slightly lesser degree than E4 ORF6 — the combined presence of E2 and E4 ORF3 did not promote increased reporter expression levels in HEK293T cells (Figure 4B). This indicates that enhanced payload expression is not solely driven by AdV genome replication but also requires a distinct molecular function uniquely executed by E4 ORF6. Probing reporter expression after co-transfection of E2 and the E4 ORF6 AXA mutant highlights that the increased payload expression requires the cellular E3 ubiquitin ligase activity, mediated by the complex of E1B-55k and E4 ORF6 (Figure 4B). This multimeric E3 ubiquitin ligase complex was shown to localize within RCs, where it promotes post-transcriptional processing and preferential export of late viral mRNA (transcribed from *de novo* synthesized genomes) from RCs to the cytoplasm while simultaneously inhibiting host mRNA export.^67,70,96^ Although the exact mechanism is still unclear, it has been hypothesized that ubiquitination-dependent relocalization of cellular export factors (e.g., the mRNA export receptor NFX1/TAP and the nuclear RNA-binding protein E1B-AP5/hRNPUL1) to RCs mediate selective export, cytoplasmic accumulation and increased translation of vector-encoded mRNA. Redistribution to RCs concurrently depletes export factors from the rest of the nuclear compartment, thereby restricting nuclear export and translation of cellular mRNA.^67,80,97,98^

Our data highlight the functionality of our designed minimal replication system. When encoded on a HC-AdV and delivered to BT-474 via transduction, it effectively drives 300-fold *cis*-replication of vector genomes and a 3.5-fold increase in MFI when compared to non-replicating HC-AdVs (Figure 5D). Transduction of HEK293T cells showed even greater genome amplification (4,000-fold) and payload expression (20-fold) (Figure 6A). Although our new vector system increased payload expression drastically, it did not fully meet the benchmark expression levels achieved by RC-AdV. We hypothesize that this discrepancy may be partially attributed to the constitutive expression of the E2 system, which lacks the replication-dependent regulation described for native AdV E2 early and E2 late promoters during the canonical genome replication of WT AdV. Perturbations in temporal expression, quantity and localization of AdV proteins have been described to jeopardize the finely tuned process of AdV replication.^99–101^ Furthermore, RC-AdVs typically employ multiple mechanisms to ensure enhanced expression of virus-encoded genes. However, our genome-replicating HC-AdV increases transgene expression primarily through genome amplification, elevated payload transcription and enhanced mRNA export. Consequently, this system lacks additional mechanisms that promote increased payload expression by mediating preferential translation of virus-encoded mRNA over cellular mRNA, as described for the virus-associated (VA) RNA_I_^102,103^ or L4-100k.^104,105^ Lastly, it is also important to consider that the enhanced payload expression levels are never directly proportional to the AdV vector copy numbers, not even upon transduction with RC-AdVs.

Our genome-replicating HC-AdV employs constitutive CMV and PGK promoters to drive expression of the E2 proteins as part of the minimal replication system (Figure 5A), upon host cell transduction. However, heterologous promoters often exhibit cell line-dependent transcriptional activities,^106,107^ potentially leading to varying E2 protein expression levels. Self-cleavage peptides additionally display different cleavage efficiencies in various cell types,^108,109^ thus the T2A peptide further contributes to divergent expression of Ad Pol within a population of transduced cells (Figure 5C) and across cell lines. Since the efficiency of our minimal replication system directly depends on sufficient expression of E2 proteins, particularly of DBP (Figure S7), varying expression levels may contribute to observed differences in genome amplification (300-fold in BT-474 vs. 4,000-fold in HEK293T cells) and enhanced payload expression (3.5-fold vs. 20-fold, respectively) upon transduction of different cell lines with our genome-replicating HC-AdV. Given the high activity of the CMV promoter in HEK293T cells^106^ correlating well with potent replication efficiency of the minimal replication system in these cells, promoters exhibiting more consistent activities across cell types may be favored for future applications of our genome-replicating HC-AdV.

The lack of morphological changes, cytopathic effects, and host cell lysis upon transduction with genome-replicating HC-AdV is indicative for reduced cytotoxicity due to the absence of intermediate and late gene expression. However, the extent to which early proteins may induce cellular toxicity, particularly in primary host cells, and whether apoptosis is effectively suppressed, remains to be elucidated. Given that AdV genomes associate with cellular chromatin during mitosis and partition into daughter cells,^110^ it will also be of great interest to explore the longevity of transgene expression throughout cell division and to determine if *in situ* AdV vector genome replication might be an integration-free mechanism of episomal transgene heredity.

The large payload capacity (>22 kb) and enhanced expression of our genome-replicating HC-AdV make it highly suitable for cancer (immuno)therapy. As we previously demonstrated, transduction with conventional HC-AdVs transforms the host cell into a “biofactory” which continuously produces therapeutic antibodies, cytokines, and/or chemokines, aiming at activating the immune system and promoting cancer cell death.^22,32,33^ While combinatorial *in situ* therapy with RC-AdVs or SC-AdVs requires combination of multiple vectors leading to cumulative toxicity effects, our genome-replicating HC-AdV provides transgene capacity to deliver multiple payloads via a single vector. Given the increased payload expression, transgene delivery via our genome-replicating HC-AdV potentially allows to reduce the required AdV vector dose and may thereby minimize the immune response and possibly prolong transgene expression for improved efficacy. Considering the large human population exhibiting pre-existing antibodies against AdV-C5, we propose deploying less prevalent AdV serotypes or chimeric capsid variants for future clinical applications in order to avoid vector inactivation mediated by preexisting neutralizing antibodies (NAbs).

Despite the advantages of genome-replicating HC-AdVs, the required expression of oncogenic E1A to promote *in situ* genome replication still remains a concern. Notably, it has been extensively demonstrated for OAdV that E1A expression can be efficiently restricted to cancer cells. This ensures that E1A is only expressed in already transformed cells, with progeny particle formation occurring exclusively in the tumor. OAdV vectors have been considered safe in multiple clinical studies.^37,38^ Consequently, tumor- and tissue-specific or regulatable expression of E1A could be a viable approach for controlling its transformation potential during cancer therapy also when using genome-replicating HC-AdVs. On the other hand, E1A-encoding AdV vector classes have also been explored for indications other than oncology. Particularly, SC-AdVs have shown promising results as a vaccination platform in several preclinical studies,^39,44,111,112^ largely attributed to the enhanced expression of the encoded antigen due to AdV genome replication.^113^

Similar to SC-AdVs, the risk of E1A-mediated transformation could be considered acceptable for certain applications of HC-AdV-mediated therapy. Our genome-replicating HC-AdV approach could be leveraged as a vaccination platform, especially in combination with our previously described retargeting adapters^114^ to specifically transduce dendritic cells (DCs), a class of professional antigen presenting cells. This strategy enables targeted transduction of DCs and to effectively boost *in situ* expression of the encoded (cancer) antigen, thereby enhancing antigen presentation and immune responses. Unlike SC-AdVs, our genome-replicating HC-AdVs offer a much larger transgene capacity, allowing for co-expression of multiple T cell stimulatory cytokines and chemokines to elicit a more potent and durable immune response against the encoded antigen.^114^ It will be of interest to investigate if our genome-replicating HC-AdVs, devoid of late genes, might be even more suitable for vaccination strategies than conventional SC-AdVs. While it has been demonstrated that oncolytic vesicular stomatitis virus (VSV) can generate tumor-reactive T cells, these leukocytes displayed signs of exhaustion due to the immunodominance of VSV antigens.^115^ This resulted in a dampened T cell response against the subdominant cancer antigens and depletion of active tumor-reactive T cells. It is therefore plausible that SC-AdVs, exhibiting high expression levels of the immunogenic late genes (e.g., hexon), may feature similarly impaired T cell responses against the encoded antigen.

It is relevant to consider current limitations of the HV-free production of genome-replicating HC-AdV vectors. Our developed vector was genetically optimized for reduced homology to the helper plasmid, however, the genome-replicating HC-AdV required the WT DBP-coding sequence (Figure S7) and the E4 promoter for efficient replication. Hence, the employed helper plasmid had to be devoid of the DBP gene and E4 cassette to reduce recombination risks during production. Consistent with findings from the production of second-generation AdV vectors, we observed that extended *trans*-complementation of multiple AdV proteins – via separate expression from vector genome and helper plasmid – significantly diminishes the vector yield. Deletion of DBP from the helper plasmid disrupts splicing branchpoints critical for efficient late transcript processing,^116^ and therefore results in suboptimal splicing and expression of AdV late proteins, and hence reduced particle formation. Additionally, helper plasmid transfection, instead of traditional HV infection,^83^ further decreases the vector yield due to a reduced number of helper genome copies and less efficient AdV protein expression, thus further exacerbating upscaling of vector titers necessary for CsCl purification. As a result, our experiments involving helper plasmid-produced HC-AdVs were performed with unpurified vector from cell lysate, complicating precise vector quantification. To further advance our genome-replicating HC-AdV technology, new HV-free methodologies for producing high-titer HC-AdV vectors are critical, including efficient helper plasmid transfection methods (e.g., electroporation), or the development of novel producer cell lines that supply most of the necessary AdV genes at the required levels in *trans*.

Further *in vivo* characterization of our genome-replicating HC-AdV requires new advanced testing models. Since murine models are not permissive for AdV replication, RC-AdVs and SC-AdVs are currently tested in immunodeficient mice with human tumor xenografts or semi-permissive species like Syrian hamsters or cotton rats.^117^ However, these models fail to fully recapitulate immune responses against the vector, AdV proteins, and antitumor effects mediated by *in situ* expression of immunotherapeutics or cancer antigens. More advanced preclinical models, such as human organoids or humanized mice,^38,117^ are needed to better evaluate the efficacy of genome-replicating HC-AdVs and the immunological consequences of their application. Nevertheless, a novel HC-AdV vector system has been introduced with significant advantages for novel applications, addressing current limitations of AdV vector-mediated therapy.

## Materials and Methods

### Cell lines

All cell lines were maintained in tissue culture flasks of 75 cm^2^ (90076, TPP, Trasadingen, Switzerland). HEK293 cells (ATCC: CRL-1573), HEK293T/17 cells (ATCC: CRL-11268), A549 cells (ATCC: CCL-185), and 911 cells^77^ were grown in complete Dulbecco’s Modified Eagle Medium (DMEM) (D6429, Sigma-Aldrich, St. Louis, MO), supplemented with 10% (v/v) heat-inactivated fetal calf serum (FCS), and 1% (v/v) penicillin/streptomycin (Sigma-Aldrich). BT-474 cells (ATCC: HTB-20) were cultured in Roswell Park Memorial Institute (RPMI) 1640 medium (21875-034, Gibco, Waltham, MA), supplemented with 10% (v/v) FCS, and 1% (v/v) penicillin/streptomycin. The HC-AdV producer cell line 116^118^ was grown in Minimal Essential Medium (MEM) (A14518-01, Gibco), supplemented with 10% (v/v) FCS, 2 mM glutamine (25030-028, Gibco), and 100 µg/ml hygromycin B (10687010, Thermo Fisher Scientific, Waltham, MA). All cell lines were maintained at 37°C and 5% CO_2_ in a humidified atmosphere, routinely tested and confirmed negative for mycoplasma contamination, and cell counting was performed using a CASY^®^ TT cell counter (OMNI Life Science, Bremen, Germany).

### Cloning of AdV expression plasmids, vDNAs, AdV mini-chromosome, minimal replication systems, HC-AdV vectors, and helper plasmids

Mammalian expression plasmids used for transfection were derived from the pcDNA3.1(+) plasmid (Thermo Fisher Scientific), and all enzymes were purchased from New England Biolabs (NEB, Ipswich, MA), unless stated otherwise. Single AdV genes were recovered either by PCR amplification from proteinase K-digested, phenol/chloroform-extracted WT AdV-C5 genomes, or by direct synthesis of cDNA (Twist Bioscience, South San Francisco, CA). AdV genes were subcloned into pcDNA3.1 using Gibson Assembly (E2621L, NEB) or restriction digestion, followed by T4 ligation (EL0011, Thermo Fisher Scientific) and expression was driven by a CMV promoter and a bovine growth hormone (bGH) polyadenylation sequence. To generate pcDNA3.1 E1, the full-length E1 TU of AdV-C5 encoding E1A, E1B-19k and E1B-55k (nt 560 – nt 4,344; GenBank AC_000008), was amplified using the following forward (5’-CTGGCTAGCGTTTAAACTTAAGCTTGGTACCACC<u>ATGAGACATATTATCTGCCACGGA</u>-3’) and reverse (5’-TGCAGAATTCTCA<u>TGGCAATCAGCTTGCTACTGA</u>-3’) primers (template hybridization sequence is underscored) and subcloned using *NheI-EcoRI*. The resulting plasmid pcDNA3.1 E1 promotes CMV-driven expression of the individual E1A isoforms and expression of E1B proteins using the endogenous E1B promoter. Plasmids encoding ΔE1,ΔE3 vDNA and ΔE1,ΔE3,ΔE4 vDNA were generated as previously described.^119^ Due to the absence of unique restriction sites, vDNAs encoding an inactive MLP were generated by assembly of multiple fragments via homologous yeast recombination (HYR).^120–122^ For this purpose, a 1131 bp gene fragment encoding multiple silencing mutations (Figure S3A) was synthesized (Twist Bioscience). This fragment included homology overlaps with vDNAs at the closest *KasI* and *XmaI* restriction site, upstream and downstream of the MLP, respectively. Three more overlapping fragments covering the remaining coding sequence of the vDNA were generated by restriction digestion, and additionally, a pRS413 *S. cerevisiae*/*E. coli* shuttle plasmid backbone^123^ was generated via PCR exhibiting overlaps to the AdV ITRs using the following forward (5’-AAGGTATATTATTGATGATGTTAATTAA<u>GAATTAATTCGATCCTGAATGGCGA</u>-3’) and reverse (5’-AAGGTATATTATTGATGATGTTAATTAA<u>TCCGCTCACAATTCCACACA</u>-3’) primer. Competent yeast cells of the *S. cerevisiae* strain VL6-48 (ATCC: MYA-3666) were prepared according to the manufacturer’s protocol using the Yeast Transformation II Kit® (T2001, Zymo Research, Irvine, CA), and 1 × 10^8^ competent yeast cells were combined with 60 fmol of each of the five DNA fragments and transformed as described by the manufacturer. After 2 h of recovery at 30°C, all cells were plated on histidine-deficient agar plates (2% (w/v) D-glucose, 0.17% (w/v) yeast nitrogen base, 0.5% (w/v) (NH_4_)_2_SO_4_, 770 mg/l CSM -His (DCS0071, Formedium, Hunstanton, UK), and 2% (w/v) bacto agar) and incubated for 3-4 days at 30°C. Single clones were used to inoculate 3 ml of histidine-deficient selection media (2% (w/v) D-glucose, 0.17% (w/v) yeast nitrogen base, 0.5% (w/v) (NH_4_)_2_SO_4_, 770 mg/l CSM -His (DCS0071, Formedium)), grown to an OD_600_ of 0.6, and yeast DNA was extracted using Yeast Plasmid MiniPrep II® (D2004, Zymo Research), as described by the supplier. Electrocompetent XL1-Blue *E. coli* cells (200159, Agilent Technologies, Santa Clara, CA) were transformed with 1/10^th^ of the eluted yeast DNA, following the manufacturer’s protocol. After outgrowth of single clones, bacterial DNA was extracted by Miniprep (27106, Qiagen, Hilden, Germany), and plasmids exhibiting correctly assembled vDNA were isolated upon analytical restriction digestion and Sanger sequencing.

An AdV mini-chromosome (amplicon) was generated by synthesis (Twist Bioscience) of a pBR322 bacterial backbone, flanked by an *EcoRI* restriction site that is directly adjacent to the 103 bp AdV ITRs,^12^ containing the core and the auxiliary origin of replication required for AdV DNA replication to occur (5’-**GAATT<u>C</u>**<u>ATCATCAATAATATACC</u>-3’; *EcoRI* restriction site denoted in bold and core origin underlined). This synthesized plasmid was then linearized and a CMV-GFP-bGH reporter cassette was subcloned in between the left and right ITR.

The semi-synthetic E2 expression system (Figure 3C) encodes a tricistronic cassette under the control of a CMV promoter, resulting in transcription of a single mRNA for DBP, pTP, and Ad Pol, and polyadenylation by the bGH termination signal. DBP translation is cap-dependent, while pTP was expressed via an IRES derived from the *encephalomyocarditis* virus (EMCV).^124^ A C-terminal GSG-linker, combined with a *Thosea asigna* virus-derived T2A self-cleavage peptide ((GSG)EGRGSLLTCGDVEENPGP),^109^ links pTP (lacking a stop codon) to Ad Pol, enabling their separate translation via ribosomal skipping. All E2 proteins encode internal FLAG-tags that were previously reported to not perturb protein functionality.^81^ Certain codons in the E2 genes were modified to facilitate subcloning by introduction or removal of unique restriction sites without altering protein sequences. The E2 expression system was assembled into a plasmid via Gibson Assembly using three synthetic DNA fragments (Twist Bioscience) and a PCR-amplified pRS413 backbone. The E4 ORF6 expression cassette encoding the endogenous E4 promoter, E4 ORF6 gene and a SV40 polyadenylation signal was subsequently cloned into the E2 expression system. To enhance DBP expression (Figure S7B) and thus replication efficiency (Figure S7C), the tricistronic cassette was split into a monocistronic (CMV promoter – DBP cDNA – bGH polyadenylation signal), and a dicistronic cassette (PGK promoter – pTP cDNA – T2A – Ad Pol cDNA – WPRE – human β-globin (hβG) polyadenylation signal) (Figure S7A, “Minimal Replication System 3”). Sequence homology of the resulting minimal replication system with HV or helper plasmids was reduced to minimize recombination risk during production by codon-optimizing pTP, Ad Pol, and E4 ORF6 and replacing homologous 5’- and 3’-untranslated regions (UTRs) (Twist Bioscience) (Figure S7A). All described modifications were performed using ligation or Gibson Assembly.

Conventional HC-AdVs, only encoding non-AdV genes, *cis*-acting elements, and stuffer DNA, were cloned as previously described.^32^ To produce HC-AdV (GFP), a GFP reporter gene was PCR amplified (5’-CACAGCTAGCTCCACC<u>ATGGTGAGCAAGGGCGAG</u>-3’, 5’- GCTTCTCGAGGCAT<u>CTATCACTTGTACAGCTCGTCCA</u>-3’). Both the PCR fragment and pUni shuttle plasmid^32^ were digested with *NheI* and *XhoI*, followed by ligation of the insert into the pUni plasmid, and thus reconstituting a functional reporter expression cassette driven by a CMV promoter and a bGH terminator. The linearized pUni shuttle plasmid was subcloned into a linearized pC4HSU backbone^16,32^ via Gibson Assembly, producing pC4HSU (GFP), which constitutes the HC-AdV genome during vector production. To generate pC4HSU (E1 + GFP), the entire E1 TU (nt 1 – nt 4,344; AC_000008) under its native E1A and E1B promoters was first subcloned into pC4HSU (GFP). For this purpose, pC4HSU (GFP) was linearized by co-digestion with *SgrAI*/*AscI*, followed by insertion of a synthetic linker (Twist Bioscience) via Gibson Assembly. This linker encoded a unique *PmeI* restriction site and, upon linearization, exhibited complementary overlaps to the E1 TU. PCR-amplified E1 TU (5’-ATGTGGCAAAAGTGACGTTT-3’, 5’-TGGCAATCAGCTTGCTACTGA-3’) was then incorporated via Gibson Assembly. This resulted in pC4HSU (E1 + GFP) which features the first 4,344 nt of WT AdV-C5. To generate pC4HSU (GPF + Minimal Replication System), pC4HSU (E1 + GFP) was digested with *PacI* and *MluI*, resulting in a 16,728 nt fragment containing the L-ITR, the packaging signal Ψ, the E1 TU, and the GFP reporter. The final version of the E2 expression system (also encoding E4 ORF6), was digested with *EcoRI* and *EcoRV*, yielding a 18,074 nt linearized fragment with homologous overlaps to *PacI*/*MluI*-linearized pC4HSU (E1 + GFP). Both fragments were assembled via homologous yeast recombination and the correct plasmid encoding pC4HSU (GFP + Minimal Replication System) was isolated upon re-transformation into the *E. coli* strain XL1-Blue.

Helper plasmids were designed as described elsewhere.^83^ First, a linear gene fragment was synthesized (Twist) encoding parts of the WT pIX cassette (nt 3,532 – nt 3,931; AC_000008), L4 TU (nt 27,082 – nt 27,336; AC_000008), and E4 ORF1 (nt 35,334 – nt 35,874; AC_000008) with unique *MfeI*-*SpeI*- and *EcoRI*-*XbaI*-restriction sites between pIX and L4, and L4 and E4 ORF1, respectively. The fragment additionally carried homologous overlaps to the pRS413 shuttle backbone, which were used to assemble both backbone and gene fragment via Gibson Assembly. Subsequently, this plasmid was linearized by *MfeI*-*SpeI*-digestion and combined via Gibson Assembly with a 23,810 bp long fragment obtained after *BstBI*-*EcoRI*-digestion of pAdEasy1.^119^ The resulting plasmid was then linearized using *EcoRI*-*XbaI* and combined via Gibson Assembly with a 5,739 bp long fragment generated from *SpeI*-*AvrII*-digested pAdEasy1. The pRS413 shuttle backbone was then replaced with a pcDNA3.1 backbone and the resulting plasmid is called pAdHelper (WT). As described by Lee *et al.*,^83^ this plasmid encodes nt 3,532 to 35,874 (AC_000008) of ΔE3 AdV-C5 but is devoid of ITRs or packaging signal Ψ and thus can virtually not be replicated or packaged in AdV capsids, resulting in efficient prevention of HV contamination. To prevent recombination during production of the genome-replicating HC-AdV encoding DBP and E4 promoter, the helper plasmid pAdHelper (ΔDBP,ΔE4) was cloned that is devoid of homologous sequences. pAdHelper (ΔDBP,ΔE4) was generated through multiple subcloning steps. To reduce homology, DBP was first deleted by removing the sequence between the L3-protease and L4-100k genes (nt 22,447 – nt 23,954; AC_000008). As a crucial late transcript splicing branchpoint is located within the N-terminal sequence of DBP,^116^ the first 78 nt of DBP were retained to not perturb late gene splicing, which otherwise impaired HC-AdV production, resulting in reduced vector yield (data not shown). Furthermore, the DBP start codon was mutated to ATA to prevent expression of truncated DBP. Next, E4 TU was deleted by removal of the AdV sequence 3’ of the fiber polyadenylation signal (nt 32,822 – nt 35,874; AC_000008), essentially resulting in ligation of the fiber cassette to the pcDNA3.1 backbone.

Sequence identity of all plasmids was confirmed via Sanger sequencing or whole plasmid sequencing using nanopore sequencing technology (Microsynth AG, Balgach, Switzerland).

### Production and purification of HC-AdV vectors using helper virus (HV)

HC-AdVs lack all AdV genes, and encode only the essential *cis*-acting elements (ITRs and Ψ), therefore they require AdV genes provided in *trans* for vector production. This is typically accomplished by co-transduction of the producer cell line 116^118^ with a HV.^125^ HC-AdVs were cloned and produced as described in detail by Brücher *et al.*^32^ The HC-AdV vector plasmid pC4HSU was *PacI*-linearized and purified by ethanol precipitation. Next, the cell line 116 (seeded 24 h prior) was transfected with the pC4HSU DNA and simultaneously co-transduced with AdV-C5 HV (lacking any capsid modifications). HC-AdV titer was sequentially increased over multiple passages by releasing the HC-AdV from the producer cells via three consecutive freeze/thaw-cycles and re-transduction of the producer cell line 116 alongside new HV. HC-AdVs were purified by cesium chloride (CsCl) gradient ultracentrifugation (250,000 × g) and dialyzed against storage buffer (20 mM HEPES pH 8.1, 150 mM NaCl, 1 mM MgCl_2_). Sterile glycerol (10% v/v) was added before snap-freezing and storage at −80°C.^32,125^

### Production of HC-AdV vectors using helper plasmid

Twenty-four hours before HC-AdV production, 12 × 10^6^ HEK293T cells were seeded in a 150 mm tissue culture dish (TPP), and pC4HSU plasmids were linearized via *PacI*-digestion and purified by ethanol precipitation. For HV-free HC-AdV production, HEK293T cells were transfected with in total 75 µg of DNA at a pC4HSU to pAdHelper ratio of 3:7 (22.5 µg pC4HSU and 52.5 µg helper plasmid). pC4HSU (GFP) and (E1 + GFP) were each co-transfected with pAdHelper (WT), while pC4HSU (GFP + Minimal Replication System) was produced using pAdHelper (ΔDBP,ΔE4). The transfection reaction was prepared in a total volume of 4 ml using serum-free Opti-MEM (31985-062, Gibco, Waltham, MA), plasmid DNA and TransIT-293 transfection reagent (2700, Mirus Bio, Madison, WI) at 1 µg DNA per 1.5 µl TransIT-293. The transfection reaction was thoroughly mixed and DNA complexation allowed to proceed for 30 min (RT), followed by gentle addition of transfection mix to HEK293T cells. At 60 h post-transfection (p.-t.), cells displayed a cytopathic effect and were harvested, pelleted (300 × g, 3 min, 4°C), resuspended in 1 ml of fresh cell culture media and subjected to three freeze-thaw cycles to release HC-AdVs. MgCl_2_ was increased to a final concentration of 5 mM, and 350 U Benzonase (E1014, Merck Millipore, Burlington, MA) were added per 1 ml cell lysate to digest free nucleic acids for 1 h at 37°C. Cell debris was removed by centrifugation (800 × g, 5 min, 4°C) and the supernatant containing the released HC-AdVs was transferred to a new sterile reaction tube. HC-AdV vector titer (transducing titer) was quantified, and the supernatant was used for transduction assays right away.

### HC-AdV vector quantification

Functionality and titers of produced vectors were determined by absorption (A_260_) and transduction assays on A549 cells. To determine the transducing titer of all AdV vectors (RC-AdVs and HV- or helper plasmid-produced HC-AdVs), 5 × 10^4^ A549 cells were seeded in a 24-well microplate (Corning, Corning, NY) 24 h prior to transduction. A549 cells were then either transduced with 3 µl of purified vector or 100 µl or producer cell lysate, containing unpurified HC-AdV. Two hours later, A549 cells were washed twice with 1 ml PBS, trypsinized, and centrifuged (800 × g, 5 min, 4°C) and the cell pellet was washed twice with PBS, followed by total DNA extraction using a Genekam DNA isolation kit (SB0072, Genekam, Germany). Transducing titers of the tested AdV vectors were quantified via multiplex-qPCR using primers and double-quenched probes binding to a distinct sequence on the HC-AdV genome (5’-TCTGCTGGTTCACAAACTGG-3’, 5’-TCCTCCCTTCTGTCCAAATG-3’, 5’-FAM-CGCCTTCTCCTGCATCCCGA-3’) or specifically hybridizing to the hexon sequence (5’-GTGATAACCGTGTGCTGGAC-3’, 5’-CAGCTTCATCCCATTCGCAA-3’, 5’-HEX-TCCGCGGCGTGCTGGACAGG-3’) (FAM = carboxyfluorescein, HEX = hexachlorofluorescein) (IDT, Coralville, IA) in order to quantify HVs, helper plasmids, and RC-AdVs, respectively, using the total DNA isolate as template. HV and helper plasmid contamination was directly determined using purified HC-AdV or unpurified HC-AdV from cell lysate as template for multiplex-qPCR. Multiplex-qPCR reactions were performed using PrimeTime Gene expression Master Mix (1055771, IDT), monitored on a Mx3000 qPCR cycler (Agilent) and analyzed as previously described.^125^

### *Trans*-replication of AdV mini-chromosome

3.6 × 10^5^ HEK293 or 1.1 × 10^5^ A549 cells were seeded in a 12-well microplate (Corning) 24 h prior to transfection. The AdV mini-chromosome was linearized by *EcoRI*-digestion and purified using QIAquick PCR purification kit (28106, Qiagen). Cells in each individual well were transfected with 1800 ng of total DNA, comprising 2 × 10^8^ copies of linearized amplicon and 2 × 10^10^ copies (33.2 fmol) of each tested plasmid, such as vDNA or AdV expression plasmid (equimolar, when used in combination). All tested AdV gene(s) were encoded on pcDNA3.1 expression plasmids. To normalize the amount of total transfected plasmid DNA to 1800 ng, empty pcDNA3.1 (circular plasmid that does not encode an expression cassette) was added. Transfection mixes were prepared in serum-free Opti-MEM (Gibco) by combining 1800 ng of total DNA and TransIT-293 transfection reagent (E1014, Mirus Bio) at a ratio of 1 µg DNA to 3 µl TransIT-293. The transfection reactions were mixed, incubated for 30 min (RT) and then gently added to the cells. Six hours p.-t., the medium was aspirated and fresh growth medium was added to the cells. Cells were harvested at 6 h and 48 h p.-t., as this provides a quantification readout pre- and post-amplicon replication. Cells were harvested by trypsinization, then centrifuged at 800 × g for 5 min (4°C) and washed twice with PBS. Total DNA was extracted from cells using DNeasy Blood & Tissue Kit (69506, Qiagen). One half (100 µl) of the DNA eluate was subjected to *DpnI*-digestion (100 U *DpnI* per digestion, 37°C, 16 h) for selective depletion of non-replicated amplicon, and subsequently total DNA was ethanol precipitated, resuspended in water, and stored at -20°C.

### *Trans*-replication of incoming, transduced HC-AdV genomes

3.6 × 10^5^ HEK293 cells were seeded in a 12-well microplate (Corning). Analogously to amplicon *trans*-replication assays, HEK293 cells were transfected 24 h later with a total of 1800 ng DNA (2 × 10^10^ copies (33.2 fmol) of each plasmid to be tested) and the remaining amount of DNA was supplemented with empty pcDNA3.1 to reach 1800 ng. The transfection mix was prepared in a total volume of 100 µl, using serum-free Opti-MEM (Gibco), 1800 ng of total DNA and TransIT-293 transfection reagent (Mirus Bio) at a ratio of 1 µg DNA to 3 µl TransIT-293. The transfection reaction was vigorously mixed, and complex formation was allowed to occur (30 min, RT) before addition of the transfection mix to the cells. Two hours p.-t., HC-AdV (GFP) virions were added to the cells for transduction at an MOI of 1. Six hours p.-t., the media was replaced with fresh growth media and cells were harvested 2 and 48 h post-transduction. Cells were washed 2× with PBS, trypsinized, and after two more PBS washes of the pellet (800 × g, 5 min, 4°C), total DNA was isolated using DNeasy Blood & Tissue Kit (Qiagen).

*Trans*-replication assays of incoming, transduced HC-AdV genomes were performed similarly in E1-non-complementing A549 cells. While 1.1 × 10^5^ A549 cells were seeded in a 12-well microplate (Corning) 24 h prior to transfection, the actual transfection mix was prepared with a total of 2500 ng DNA. The transfection mix comprised 3.6 × 10^10^ copies (59.8 fmol) of each plasmid tested for replication. The transfection mix was prepared in a total volume of 100 µl, using serum-free Opti-MEM (Gibco), 2500 ng of total DNA (supplemented to reach this amount with empty pcDNA3.1) and TransIT-293 transfection reagent (Mirus Bio) at a ratio of 1 µg DNA to 3 µl TransIT-293. After thorough mixing and incubation for 30 min (RT), the transfection mix was subsequently added to A549 cells, and it was proceeded with transduction and harvest as described above.

### Quantitative polymerase chain reaction analysis

Total DNA extracted from replication assays was used for qPCR analysis on a Mx300 (Agilent) instrument to quantify AdV vector copy numbers. qPCR analysis was performed in a 96-well qPCR plate (72.1979.010, Sarstedt, Nümbrecht, Germany) with a total reaction volume of 20 µl per well, including 5 µl of total DNA, 10 µl of 2× qPCR Mastermix (A25742, Thermo Fisher Scientific), forward and reverse primers (500 nM final concentration each), and water. The qPCR cycling protocol included a 2-minute polymerase activation at 50°C, followed by 2 minutes at 95°C, and 40 amplification cycles of 15 seconds denaturation (95°C) and 1 minute of annealing and extension (60°C). A dissociation step was performed by slowly increasing (1.6°C per s) the temperature to 95°C (15 s), slowly decreasing (1.6°C per s) to 60°C (1 min) and a final increase to 95°C for 15 s (increase of 0.15°C per s). Absolute quantification was achieved by referencing obtained Ct values to those from a standard curve (1–10^7^ copies/µl). HC-AdV (GFP) vector copy numbers were quantified using GFP primers (5’-CCGACAAGCAGAAGAACGGC-3’, 5’-GTGATCGCGCTTCTCGTTGG-3’) and normalized to cellular DNA using GAPDH primers (5’-AATTCCATGGCACCGTCAAG-3’, 5’-ATCGCCCCACTTGATTTTGG-3’).

### Payload expression assays and analysis via flow cytometry

To drive increased fluorescent reporter expression through replication of an incoming, transduced HC-AdV genome, potent genome replication is essential, hence requiring high expression levels of the replication machinery. Instead of standard HEK293 cells, HEK293T cells were thus used to enhance expression of the replication machinery due to their high transfectability and expression of the SV40 large T antigen, which supports episomal replication of plasmids containing the SV40 origin, further boosting the supply of the replication system. Prior to performing *trans*-replication assays in HEK293T cells, all plasmid backbones were thus removed by *PacI*-digestion and replaced via Gibson Assembly with PCR-amplified pcDNA3.1 backbones (5’-AAGGTATATTATTGATGATGTTAATTAA<u>TCTGAGGCGGAAAGAACCAG</u>-3’, 5’-AAGGTATATTATTGATGATGTTAATTAA<u>ACTTTTCGGGGAAATGTGCG</u>-3’) encoding the SV40 origin of replication.

To quantify changes in payload expression, 3 × 10^5^ HEK293T were seeded per well in a 6-well cell culture plate (Corning) 24 h prior to transfection. HEK293T cells were transfected with 5000 ng total DNA, comprising 1 × 10^11^ copies (166.1 fmol) of each tested plasmid and the amount of total DNA was normalized to 5000 ng using empty pcDNA3.1. Transfection reactions (250 µl total) were prepared in serum-free Opti-MEM (Gibco) by combining plasmid DNA and TransIT-293 transfection reagent (Mirus Bio) at a ratio of 1 µg DNA to 1.5 µl TransIT-293, followed by thorough mixing and incubation for 30 min (RT). Subsequently, the transfection reactions were added to the cells. Two hours p.-t., cells were transduced with the indicated HC-AdV (e.g., pC4HSU (GFP)) at a designated MOI and at 6 h p.-t., culture media was replaced with fresh growth media. Cells were harvested 72 h post-transduction and washed twice (800 × g, 5 min, 4°C) with PBS. 20% of the cells were used for total DNA isolation using a DNeasy Blood & Tissue Kit (Qiagen) and subjected to qPCR analysis, while the remaining 80% of the cells were prepared for flow cytometric analysis by resuspension in fixation buffer (PBS containing 4% (w/v) paraformaldehyde (PFA)) and fixed for 15 min at RT. Subsequently, cells were centrifuged (800 × g, 5 min, 4°C), washed once with FACS buffer (PBS containing 1 % (w/v) BSA and 0.1% (w/v) NaN_3_) and ultimately resuspended in 200 µl FACS buffer and stored at 4°C until analysis at a BD FACSymphony 5L flow cytometer (BD Biosciences, Franklin Lakes, NJ) using the high-throughput sampler. Flow cytometric analysis was performed using FlowJo™ software (BD Biosciences).

### Western blot analysis

For experiments requiring transfection, 1.5 × 10^6^ HEK293 cells were seeded into 6-well microplates (Corning) 24 h prior to transfection. Then, cells were transfected with 2500 ng of respective pcDNA-based expression plasmids using TransIT-293 (Mirus Bio) according to the manufacturer’s guidelines. For transduction experiments, 3.6 × 10^5^ BT-474 cells were seeded 24 h prior to transduction with the indicated HC-AdV at the specified MOI. Thirty-six (or as indicated) hours after transfection or transduction, cells were washed once in PBS and harvested by trypsinization. After two further washes (800 × g, 5 min, 4°C) with ice-cold PBS, cells were lysed in ice-cold lysis buffer (1% (v/v) Triton X-100, 40 mM HEPES pH 7.4, 10 mM β-glycerol phosphate, 10 mM pyrophosphate, 2.5 mM MgCl_2_, and 1 tablet of EDTA-free protease inhibitor (A32955, Thermo Fisher Scientific) per 10 mL buffer). Cell lysates were clarified by centrifugation at 21,000 × g at 4°C for 10 min, and total protein concentration was determined by BCA assays (23225, Thermo Fisher Scientific), followed by addition of 6× Laemmli Buffer to the protein extract. Protein extracts were boiled for 10 min (95°C) and normalized amounts of protein were resolved by 4-12% SDS-PAGE (NW04122, Thermo Fisher Scientific) and subsequently immunoblotted by standard wet transfer (100 V, 2 h). Nitrocellulose membranes (10401196, Whatman, Maidstone, UK) were blocked for 1 h in TBS-T (20 mM Tris-HCl, 150 mM NaCl, 0.1% (v/v) Tween 20, pH 7.4) + 5% (w/v) milk powder and membranes were incubated over-night on a roller-shaker (4°C) in TBS-T + 5% milk powder containing the corresponding primary antibody (α-FLAG M2 (1:1,000 dilution, F3165, Sigma-Aldrich); α-β-Actin (1:3,000 dilution, A3853, Sigma-Aldrich); α-GFP (1:3,000 dilution, 600-401-215, Rockland Immunochemicals, Limerick, PA); α-AdV-C5 L5-Fiber (1:10,000 dilution, ab76551, Abcam, Cambridge, UK); α-AdV-C5 E1A (1:5,000 dilution, in-house produced from murine hybridoma clone M73, purified supernatant); α-AdV-C5 DBP (1:10,000 dilution, in-house produced from murine hybridoma clone B6-8, purified supernatant); α-AdV-C5 L1-52/55k (1:1,000 dilution, in-house produced antiserum); α-AdV-C5 L4-100k (1:1,000 dilution, in-house produced antiserum)). Membranes were washed 3× for 5 min with TBS-T on a roller-shaker and then incubated with respective horseradish peroxidase (HRP)-coupled secondary antibody (α-mouse IgG (1:10,000 dilution, 31438, Thermo Fisher Scientific); α-rabbit IgG (1:5,000 dilution, 7074, Cell Signaling Technology, Danvers, MA)), diluted in TBS-T for 1 h (RT) on a roller-shaker. After binding of the secondary antibody, membranes were washed again 3× with TBS-T for 5 min each, ECL substrate solution (WBKLS0050, Merck Millipore) was added and protein bands were visualized using a FusionFX Imaging System (Vilber, Collégien, France).

## Data and code availability

The data presented in this study are available upon reasonable request to the corresponding author A.P.

## Acknowledgments

We thank Dr. Philip Ng (Baylor College of Medicine) for kindly providing the cell line 116 and the HV AdNG163R-2, and Prof. Urs Greber (University of Zurich) for sharing the producer cell line 911 with us. Furthermore, we are thankful to Prof. Patrick Hearing (Renaissance School of Medicine at Stony Brook University) for sharing in-house produced α-AdV-C5 L1-52/55k and α-AdV-C5 L4-100k antibodies as well as the murine hybridoma cell lines M73 (Harlow Lab) and B6-8 (Levine Lab) with us. We are very grateful for valuable scientific discussion with Prof. Urs Greber, Prof. Anja Ehrhardt (University of Witten/Herdecke), and Prof. Patrick Hearing. We acknowledge Dr. Dominik Brücher for providing materials and reagents, and we would like to thank Dr. K. Patricia Hartmann and Dr. Davor Nestić for helpful discussions and critical reading of the manuscript. We acknowledge the Flow Cytometry Facility of the University of Zurich for training of users and maintenance of the instruments. We would like to thank the following funding agencies for supporting this work: the University of Zurich (Candoc Grant FK-20-031 to J.K.) and the Swiss National Science Foundation (Sinergia Grant CRSII5_170929 to A.P.). The figures were partially generated with BioRender.com.

## Author contributions

Conceptualization: J.K., and A.P.; Methodology: J.K.; Investigation: J.K., F.W., P.C.F.; Formal Analysis: J.K.; Resources: J.K.; Validation: J.K.; Project Administration: J.K.; Visualization: J.K.; Supervision: A.P.; Writing – Original Draft: J.K.; Writing – Review & Editing: J.K., F.W., P.C.F., and A.P.; Funding Acquisition: J.K., and A.P.

