## Supplementary Materials for "Genome-replicating HC-AdV – a novel high-capacity adenoviral vector class featuring enhanced *in situ* payload expression"

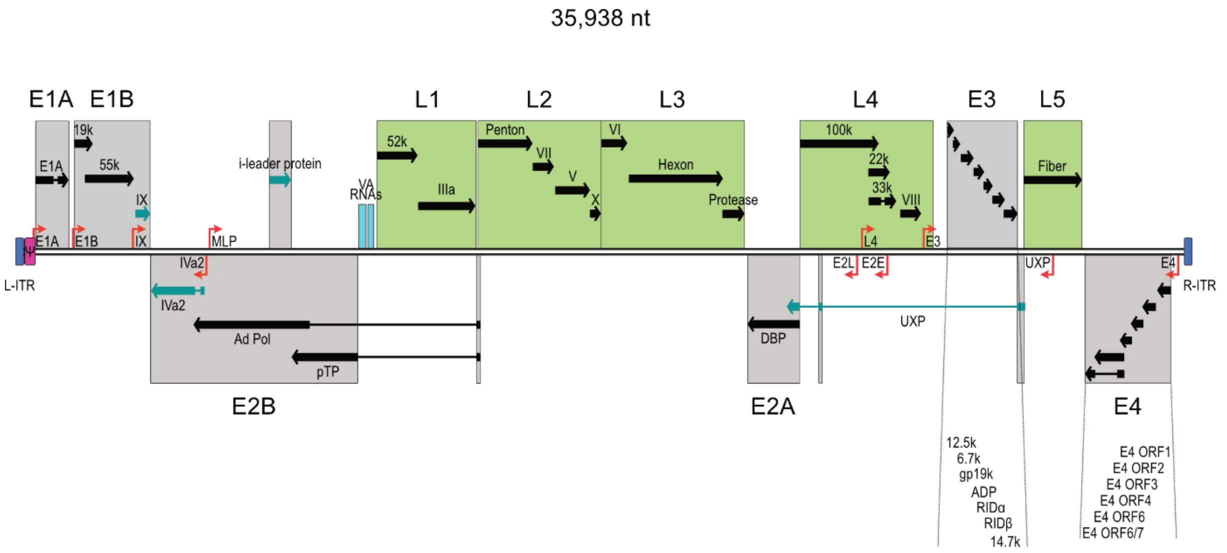


**Figure S1. Schematic illustration of the genetic architecture of the human wild-type AdV-C5 genome.** The double-stranded DNA genome is displayed as a pair of black lines, left and right inverted terminal repeats (ITRs) are shown in blue and the packaging signal Ψ in magenta. Transcriptional units (TUs) are depicted in gray (early TU 1-4 and intermediate TUs) and green (late TU 1-5) boxes with individual transcripts shown as black arrows with the thick bars denoting open reading frames (ORFs) and thin bars denoting introns and untranslated regions (UTRs). Intermediate transcripts are displayed in teal and adenoviral virus-associated RNA (VA RNA) I and II are illustrated in cyan. Transcription start sites (TSS) of TUs or individual genes are represented by red arrows.

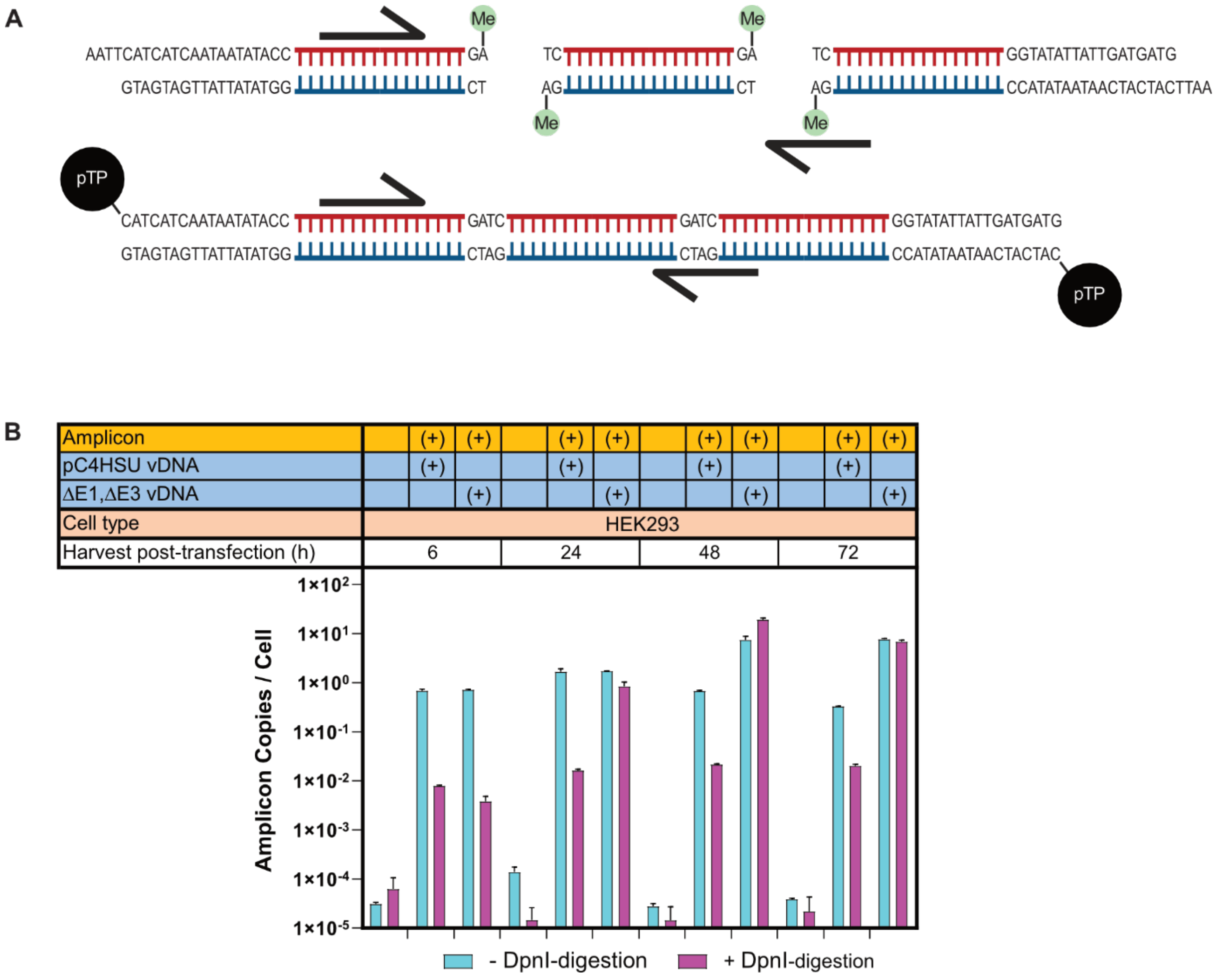


**Figure S2. *DpnI*-digestion allows selective quantification of *de novo* synthesized amplicons via qPCR.** (A) Schematic depiction of transfected amplicon (upper panel) and *de novo* replicated amplicon (lower panel). Only the transfected amplicon is selectively depleted, as successful *DpnI*-digestion requires methylation of recognition site (green), mediated by the bacterial Dam methylase during production of DNA molecules in *E. coli*. The *de novo* synthesized amplicon is replicated in mammalian cells, not methylated and thus resistant to *DpnI* digestion and provides a valid template for qPCR analysis. qPCR primers (black) can bind to the correct DNA template for selective quantification of amplicon copy numbers. (B) *Trans*-replication leads to resistance of the amplicon towards *DpnI*-digestion, as determined by qPCR analysis. Amplicon copies per cell were determined after co-transfection with plasmids reconstituting a functional (e.g., ΔE1,ΔE3 vDNA) or non-functional replication machinery (pC4HSU vDNA). Quantification of amplicons was performed 6, 24, 48 and 72 h post-transfection via qPCR, after total DNA was left untreated (cyan bars) or *DpnI*-digested (magenta bars). Bar graphs represent mean amplicon copy numbers per cell ± SD, n = 3.

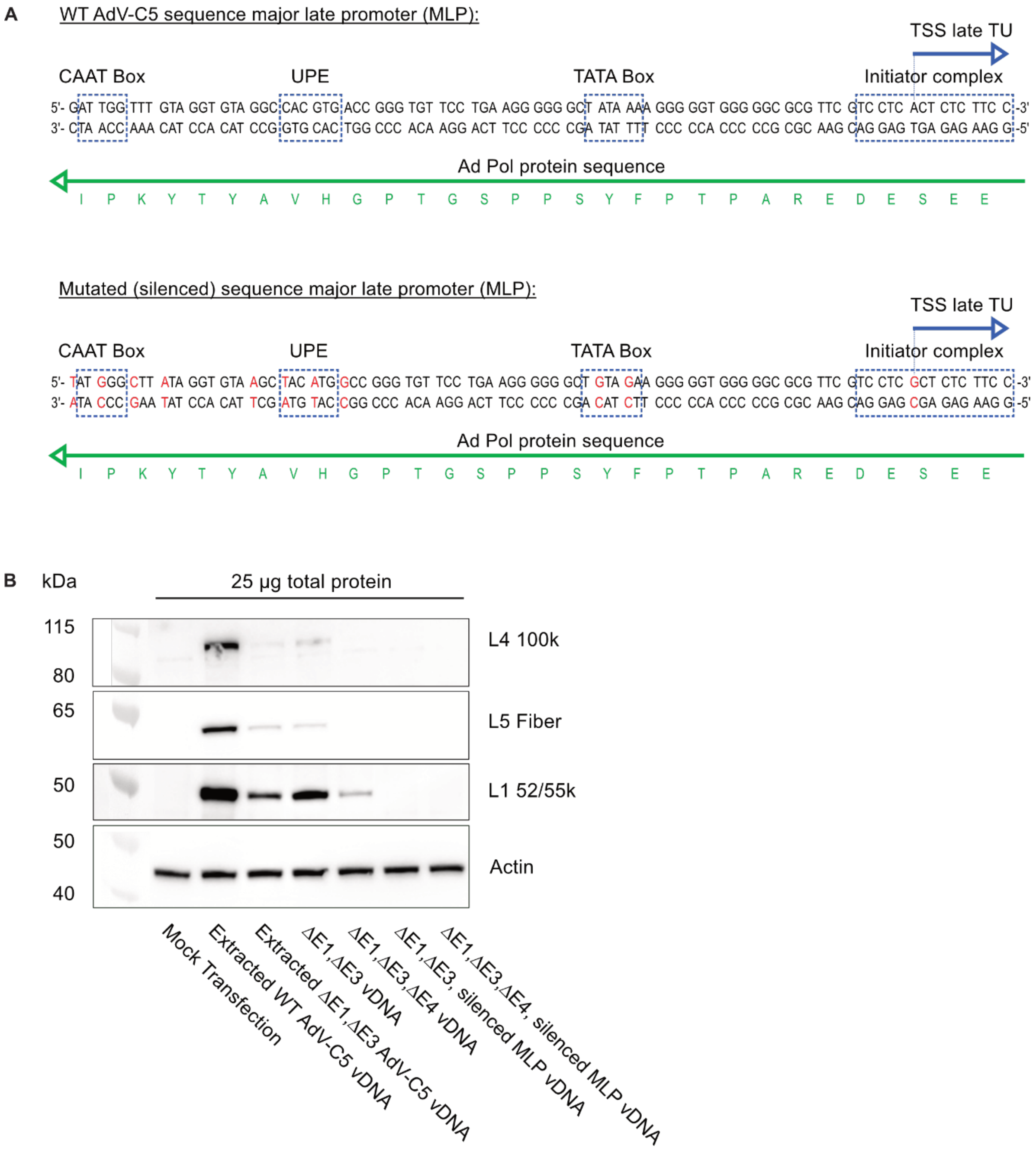


**Figure S3. Silencing of major late promoter (MLP).** (A) Sequence of the AdV-C5 MLP. The wild-type nucleotide sequences (upper panel) of transcription factor binding sites (CAAT Box, UPE, and TATA Box), binding sequence of the initiator complex, and transcription start site (TSS) of the major late transcript are denoted on the sense strand with a blue dashed box. The nucleotide sequence of the Ad Pol is encoded on the antisense strand and the corresponding amino acid sequence is displayed below in green. The mutated nucleotide sequence of the silenced MLP is illustrated in the lower panel. Multiple disrupting point mutations in transcription factor and initiator complex binding sites are highlighted in red. Point mutations preserved the protein-coding sequence of the Ad Pol, encoded on the antisense strand, as shown by the Ad Pol amino acid sequence (green). (B) Representative western blot analysis and immunostaining of three late gene products from the L1, L4, and L5 TU, 72 h post-transfection of HEK293 cells with different vDNAs. Cells were transfected with WT AdV-C5 vDNA and ΔE1,ΔE3 AdV-C5 vDNA extracted from functional purified virions via proteinase K digestion and phenol/chloroform-extraction (lane 2 and 3). Additional transfections included ΔE1,ΔE3 vDNA (lane 4), ΔE1,ΔE3,ΔE4 vDNA (lane 5), ΔE1,ΔE3, silenced MLP vDNA (lane 6), and ΔE1,ΔE3,ΔE4, silenced MLP vDNA (lane 7), all produced as plasmids in *E. coli*.

| Amplicon | (+) | (+) | (+) | (+) | (+) | (+) | (+) | (+) | (+) | (+) | (+) |
|---|---|---|---|---|---|---|---|---|---|---|---|
| ΔE1,ΔE3 vDNA | (+) | | | | | | | | | | |
| ΔE1,ΔE3,ΔE4 vDNA | | (+) | (+) | (+) | (+) | (+) | (+) | (+) | | | |
| ΔE1,ΔE3, silenced MLP vDNA | | | | | | | | | (+) | | |
| ΔE1,ΔE3,ΔE4, silenced MLP vDNA | | | | | | | | | | (+) | (+) |
| pcDNA3.1 E4 ORF1 | | | (+) | | | | | | | | |
| pcDNA3.1 E4 ORF2 | | | | (+) | | | | | | | |
| pcDNA3.1 E4 ORF3 | | | | | (+) | | | | | | |
| pcDNA3.1 E4 ORF4 | | | | | | (+) | | | | | |
| pcDNA3.1 E4 ORF6 | | | | | | | (+) | | | | (+) |
| pcDNA3.1 E4 ORF6/7 | | | | | | | | (+) | | | |
| Cell type | 911 | | | | | | | | | | |

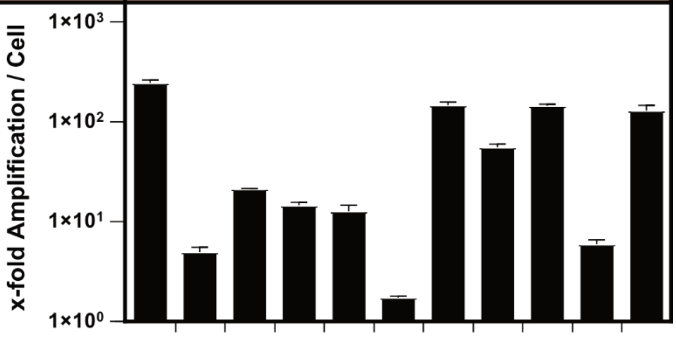


**Figure S4. Characterization of E4 and late genes via *trans*-replication of amplicon in the cell line 911.** Quantification of x-fold *trans*-amplification per cell via qPCR, 48 h after co-transfection of 911 cells with amplicon (orange), ΔE1,ΔE3 vDNA, ΔE1,ΔE3,ΔE4 vDNA, ΔE1,ΔE3, silenced MLP vDNA, and ΔE1,ΔE3,ΔE4, silenced MLP vDNA (dark blue). Co-transfection of amplicon and ΔE1,ΔE3,ΔE4 vDNA was additionally *trans*-complemented with all different gene products form the E4 TU (light blue). Bar graphs represent x-fold amplification ± SD, n = 3.

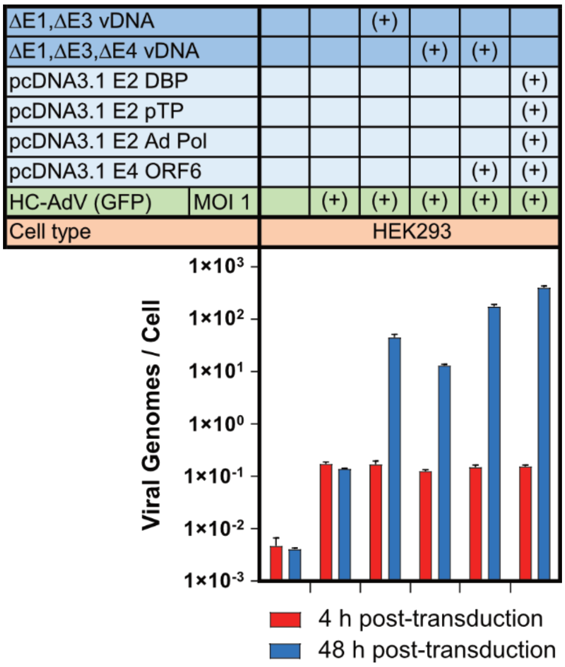


**Figure S5. *Trans*-replication of incoming, transduced HC-AdV genomes upon pre-transfection of HEK293 cells with replication systems.** Quantification of incoming, transduced HC-AdV genome replication via qPCR upon pre-transfection of HEK293 cells with AdV replication machineries (dark blue) or individual AdV genes (light blue). HC-AdV copy numbers were quantified 4 h (red bars) and 48 h (blue bars) after transduction. Bar graphs represent mean HC-AdV copy numbers per cell ± SD, n = 3.

| | | | | | | | | | | | | | | | | | | | | |
|---|---|---|---|---|---|---|---|---|---|---|---|---|---|---|---|---|---|---|---|---|
| ΔE1,ΔE3 vDNA | (+) | | | | | | | | | | | | | | | | | | | |
| E2 Expression System | | | (+) | (+) | (+) | (+) | (+) | (+) | (+) | (+) | (+) | (+) | (+) | (+) | (+) | (+) | (+) | (+) | (+) | (+) |
| pcDNA3.1 E1 | (+) | (+) | (+) | (+) | (+) | (+) | (+) | (+) | (+) | (+) | (+) | (+) | (+) | (+) | (+) | (+) | (+) | (+) | (+) | (+) |
| pcDNA3.1 E2 DBP | | (+) | | | | | | | | | | | | | | | | | | |
| pcDNA3.1 E2 pTP | | (+) | | | | | | | | | | | | | | | | | | |
| pcDNA3.1 E2 Ad Pol | | (+) | | | | | | | | | | | | | | | | | | |
| pcDNA3.1 E4 ORF6 | | (+) | (+) | (+) | (+) | (+) | (+) | (+) | (+) | (+) | (+) | (+) | (+) | (+) | (+) | (+) | (+) | (+) | (+) | (+) |
| pcDNA3.1 IVa2 | | | | (+) | | | | | | | | | | | (+) | | | (+) | (+) | (+) |
| pcDNA3.1 IX | | | | | (+) | | | | | | | | | | (+) | | | | | (+) |
| pcDNA3.1 UXP | | | | | | (+) | | | | | | | | | (+) | | | | (+) | (+) |
| pcDNA3.1 i-leader protein | | | | | | | (+) | | | | | | | | (+) | | | | | (+) |
| pcDNA3.1 L1 pIIIa | | | | | | | | (+) | | | | | | | | (+) | | | | |
| pcDNA3.1 L1 p52k | | | | | | | | | (+) | | | | | | | (+) | | | | |
| pcDNA3.1 L2 pVII | | | | | | | | | | (+) | | | | | | | | | | |
| pcDNA3.1 L4 100k | | | | | | | | | | | (+) | | | | | | (+) | | | |
| pcDNA3.1 L4 22k | | | | | | | | | | | | (+) | | | | | (+) | (+) | (+) | (+) |
| pcDNA3.1 L4 33k | | | | | | | | | | | | | (+) | | | | (+) | (+) | (+) | (+) |
| pcDNA3.1 L4 pVIII | | | | | | | | | | | | | | (+) | | | (+) | | | |
| HC-AdV (GFP) MOI 1 | (+) | (+) | (+) | (+) | (+) | (+) | (+) | (+) | (+) | (+) | (+) | (+) | (+) | (+) | (+) | (+) | (+) | (+) | (+) | (+) |
| Cell type | A549 | | | | | | | | | | | | | | | | | | | |

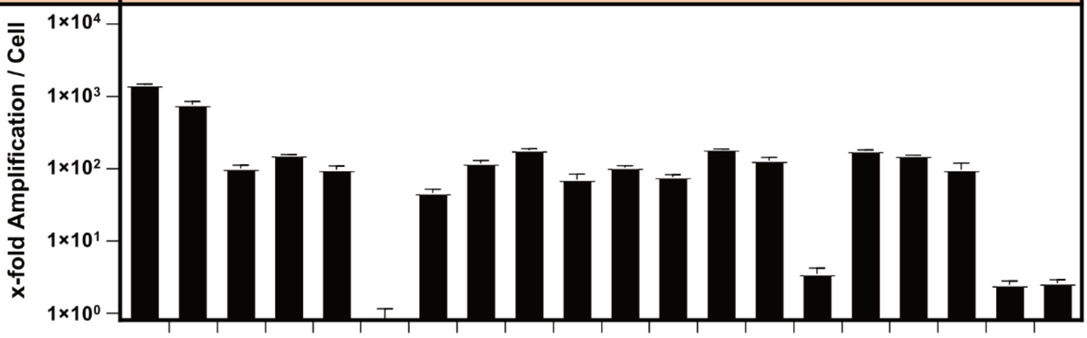


**Figure S6. *Trans*-replication of incoming, transduced HC-AdV genomes upon pre-transfection of A549 cells with replication systems and intermediate and/or late genes.** Quantification of x-fold amplification of incoming, transduced HC-AdV genomes via qPCR, as analyzed 48 h post-transduction of E1-non complementing A549 cells with different replication systems (dark blue). The E2 expression system was co-transfected with plasmids encoding E1, E4 ORF6, and individual or combined expression plasmids encoding relevant intermediate genes (IVa2, IX, UXP, and i-leader protein) or late genes (L1 pIIIa, L1 p52k, L2 pVII, L4 100k, L4 22k, L4 33k, and L4 pVIII) (light blue). Bar graphs represent x-fold amplification ± SD, n = 3.

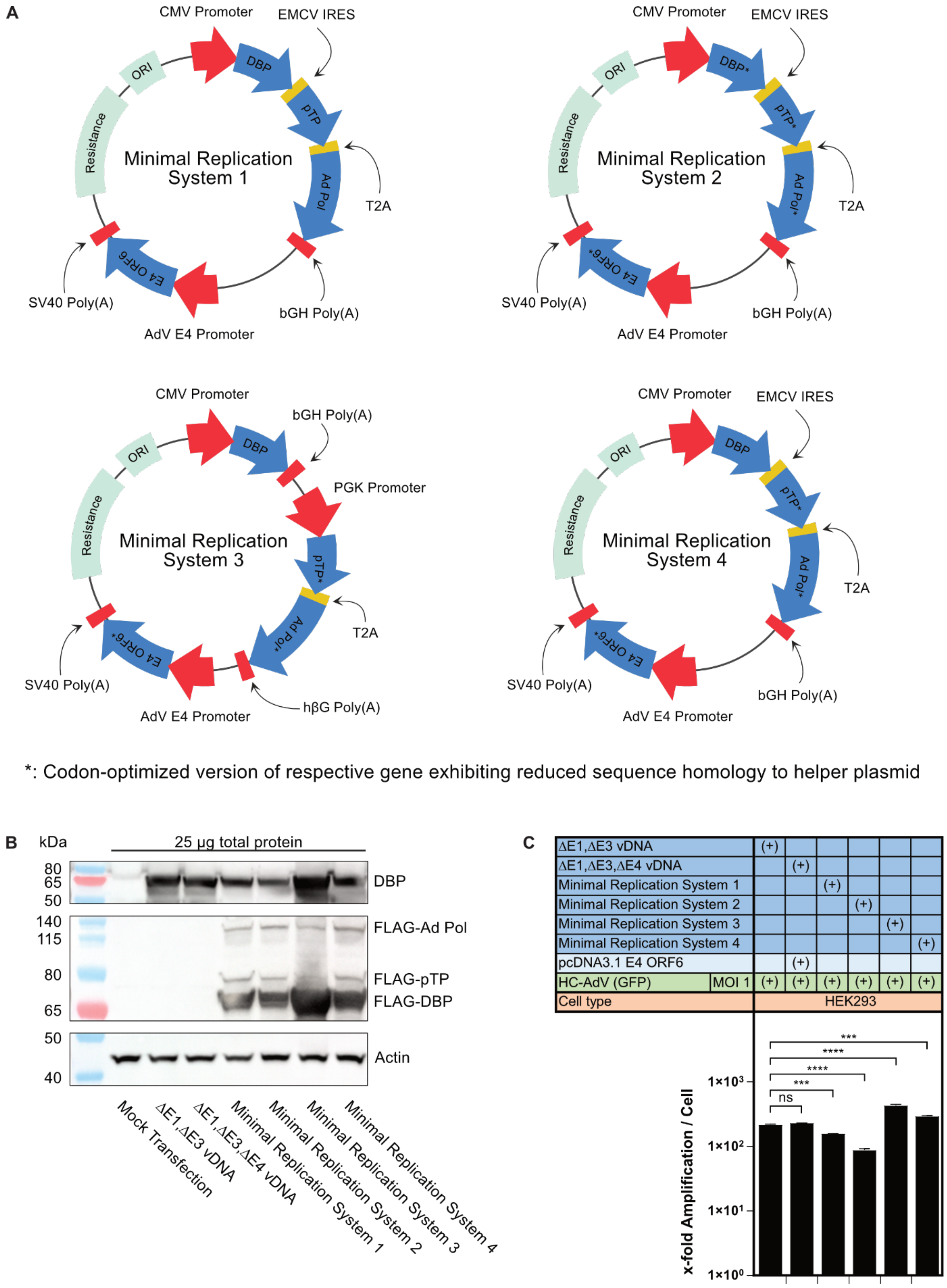


**Figure S7. Development and characterization of different minimal replication systems.** (A) Schematic depiction of engineered E2 expression systems additionally encoding E4 ORF6 in a separate expression cassette. In combination with E1 (supplied in *trans* or *cis*), these constructs reconstitute a minimal AdV replication system. Minimal Replication System 1 features a tricistronic cassette expressing all WT E2 proteins under a CMV promoter with a bovine growth hormone (bGH) transcription termination sequence, and a separate cassette expressing E4 ORF6 (WT) driven by the endogenous AdV E4 promoter and an SV40 polyadenylation signal. Minimal replication system 2 exhibits an identical configuration of expression cassettes as system 1, but with codon-optimized sequences for the three E2 genes and E4 ORF6 to reduce sequence homology with helper plasmids (denoted by asterisk). The modified nucleotide sequences only contain silent mutations and thus preserve the WT protein-coding sequences. Minimal Replication Systems 3 and 4 both encode WT DBP alongside codon-optimized pTP, Ad Pol, and E4 ORF6. In system 3, WT DBP is supplied from a separate expression cassette (using a CMV promoter and a bGH polyadenylation signal) while codon-optimized pTP and Ad Pol are expressed from a dicistronic cassette using a PGK promoter, a T2A self-cleavage sequence, and a human β-globin termination (hβG) signal. Minimal replication system 4 is identical to system 2 except that it expresses WT DBP instead of the codon-optimized version. (B) Representative western blot analysis and immunostaining of DBP and FLAG-tagged E2 proteins, 36 h post-transfection of HEK293 cells with ΔE1,ΔE3 vDNA, ΔE1,ΔE3,ΔE4 vDNA, or minimal replication system 1-4. Detection of FLAG-tag allows evaluation of E2 protein expression across minimal replication systems, while DBP immunostaining enables direct comparison of DBP expression levels from replication systems to tag-less vDNAs. β-actin was immunostained as a loading control. (C) Quantification of incoming, transduced HC-AdV genome replication via qPCR upon pre-transfection of HEK293 cells with different vDNAs or minimal replication systems (dark blue). The ΔE1,ΔE3,ΔE4 vDNA was co-transfected with plasmids encoding E4 ORF6 (light blue). Genome amplification was quantified 48 h post-transduction and is presented as x-fold replication. Bar graphs represent x-fold amplification ± SD, n = 3. Statistical significance was determined by one-way ANOVA with Dunnett's test for multiple comparisons to the control group of transfected ΔE1,ΔE3 vDNA. Not significant (ns) $P > 0.05$; *$P \leq 0.05$; **$P \leq 0.01$; ***$P \leq 0.001$; ****$P \leq 0.0001$.

| | | | | | | | | | | | | | |
|---|---|---|---|---|---|---|---|---|---|---|---|---|---|
| ΔE1,ΔE3 vDNA | (+) | | | | | | | | | | | | |
| E2 Expression System | | | | (+) | (+) | (+) | (+) | (+) | | | | | |
| E2 Expression System (Ad Pol KO) | | | | | | | | | (+) | (+) | | | |
| Minimal Replication System (CMV-E2 + PGK-E4 ORF6) | | | | | | | | | | | (+) | | |
| Minimal Replication System (CMV-E2 + SV40-E4 ORF6) | | | | | | | | | | | | (+) | |
| Minimal Replication System (CMV-E2 + E4-E4 ORF6) | | | | | | | | | | | | | (+) |
| pcDNA3.1 CMV-E2 DBP | | (+) | (+) | | | | | | | | | | |
| pcDNA3.1 CMV-E2 pTP | | (+) | (+) | | | | | | | | | | |
| pcDNA3.1 CMV-E2 Ad Pol | | (+) | (+) | | | | | | | | | | |
| pcDNA3.1 CMV-E4 ORF6 | | | (+) | | (+) | | | | | (+) | | | |
| pcDNA3.1 PGK-E4 ORF6 | | | | | | (+) | | | | | | | |
| pcDNA3.1 SV40-E4 ORF6 | | | | | | | (+) | | | | | | |
| pcDNA3.1 E4-E4 ORF6 | | | | | | | | (+) | | | | | |
| HC-AdV (GFP) MOI 1 | (+) | (+) | (+) | (+) | (+) | (+) | (+) | (+) | (+) | (+) | (+) | (+) | (+) |
| Cell type | HEK293 | | | | | | | | | | | | |

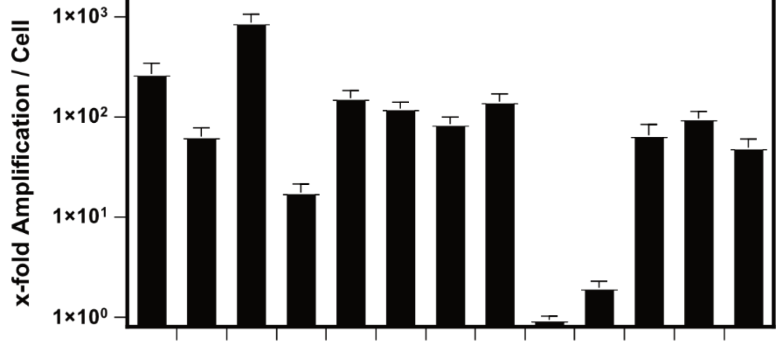


**Figure S8. Effect of different promoters driving E4 ORF6 expression in conjunction with CMV-driven E2 expression system on *trans*-replication of incoming, transduced HC-AdV genomes in HEK293 cells.** Quantification of x-fold HC-AdV genome replication via qPCR upon pre-transfection of HEK293 cells with AdV replication machineries (dark blue) or individual AdV genes (light blue). The CMV-driven E2 expression system was *trans*-complemented with individual plasmids expressing E4 ORF6 under the control of the CMV, PGK, SV40 or endogenous AdV E4 promoter. Alternatively, the plasmid encoding the CMV-driven E2 expression system was modified to additionally encode an E4 ORF6 expression cassette driven by either PGK, SV40, or AdV E4 promoter and pre-transfected for *trans*-replication of incoming HC-AdV genomes. Bar graphs represent x-fold amplification ± SD, n = 3.
